# AMP-Kinase mediates regulation of glomerular volume and podocyte survival

**DOI:** 10.1101/2021.05.21.445180

**Authors:** Khadija Banu, Qisheng Lin, John M. Basgen, Marina Planoutene, Chengguo Wei, Anand C. Reghuvaran, Felipe Garzon, Aitor Garcia, Nicholas Chun, Arun Cumpelik, Hongmei Shi, Andrew Santaneusio, Weijia Zhang, Bhaskar Das, Fadi Salem, Li Li, Lloyd G Cantley, Shuta Ishibe, Lewis Kaufman, Kevin V. Lemley, Zhaohui Ni, John C. He, Barbara Murphy, Madhav C. Menon

**Author notes:** Equal contribution (Khadija Banu, Qisheng Lin). **Correspondence:** Madhav C Menon, MBBS, MD FACP, Nephrology, Department of Medicine, Yale University School of Medicine, 300 Cedar Street, 255A, New Haven, CT 06519, USA, or.

## Abstract

We reported that Shroom3 knockdown, via Fyn inhibition, induced albuminuria with foot process effacement (FPE) without glomerulosclerosis (FSGS) or podocytopenia. Interestingly, knockdown mice had reduced podocyte volumes. Human minimal change disease, where podocyte Fyn inactivation was reported, also showed lower glomerular volumes than FSGS. We hypothesized that lower glomerular volume prevented the progression to podocytopenia. To test this hypothesis, we utilized unilateral- and 5/6^th^ nephrectomy models in Shroom3 knockdown mice. Knockdown mice exhibited lower glomerular volume, and less glomerular and podocyte hypertrophy after nephrectomy. FYN-knockdown podocytes had similar reductions in podocyte volume, implying Fyn was downstream of Shroom3. Using SHROOM3- or FYN-knockdown, we confirmed reduced podocyte protein content, along with significantly increased phosphorylated AMP-kinase, a negative regulator of anabolism. AMP-Kinase activation resulted from increased cytoplasmic redistribution of LKB1 in podocytes. Inhibition of AMP-Kinase abolished the reduction in glomerular volume and induced podocytopenia in mice with FPE, suggesting a protective role for AMP-Kinase activation. In agreement with this, treatment of glomerular injury models with AMP-Kinase activators restricted glomerular volume, podocytopenia and progression to FSGS. In summary, we demonstrate the important role of AMP-Kinase in glomerular volume regulation and podocyte survival. Our data suggest that AMP-Kinase activation adaptively regulates glomerular volume to prevent podocytopenia in the context of podocyte injury.

## Introduction

Podocytes are specialized epithelial cells on the urinary side of the glomerular filtration barrier with interdigitating cellular extensions or foot processes. Podocyte actin cytoskeletal disorganization is near universal with nephrotic syndrome (NS) and visualized as foot process effacement (FPE) (1).

Focal segmental glomerulosclerosis (FSGS) causes proteinuria and NS where podocytes show diffuse FPE associated with podocyte loss, glomerulosclerosis and progressive renal failure (2), while minimal change disease (MCD), in spite of diffuse FPE, shows no podocytopenia and low rates of disease progression. By morphometry, FSGS has been associated with larger glomerular volumes (Vglom) and podocyte hypertrophy (3, 4), and with adverse clinical outcomes (3, 5), (6, 7). After unilateral nephrectomy, increases of Vglom in remnant kidneys disproportionate to podocyte hypertrophy, also promoted FSGS and podocytopenia (8). By contrast, the specific significance of Vglom in MCD is unknown. Furthermore, whether regulating Vglom in the context of podocyte FPE and injury could offer benefit to podocyte survival, needs to be examined. A mechanism for regulating glomerular volume would be of therapeutic interest if it mitigated podocytopenia and progression to FSGS following glomerular injury and podocyte FPE.

Our group, and others, have serially reported on the dichotomous roles of Shroom3 in tubular cells and podocytes in kidney disease (9-12). Using an inducible knockdown model, in young mice (8-12 weeks old), we recently reported that glomerular Shroom3 knockdown caused proteinuria with diffuse FPE (10). We uncovered a novel protein-protein interaction between Shroom3 and Fyn that regulated Fyn activation. In podocytes in this model, inhibited Nphs1 phosphorylation downstream of Fyn inactivation, caused actin cytoskeletal disorganization and FPE. Fyn is a non-receptor tyrosine kinase and regulates multiple signaling cascades via phosphorylation of tyrosine residues (13). Interestingly, podocyte Fyn inactivation was also specifically associated with human MCD where diffuse FPE without podocytopenia is seen (14, 15). Analogously, in young mice with Fyn inactivation following Shroom3 knockdown, glomeruli showed no podocytopenia or FSGS in spite of diffuse FPE, an “MCD-like” pathology. We also observed that podocyte volume was reduced by Shroom3 knockdown both *in vitro* and *in vivo* (10).

Based on this surprising morphometric finding, we first examined a cohort of human NS cases from the Neptune consortium. We identified significantly reduced Vglom in MCD vs FSGS, suggesting maintained Vglom regulation in MCD cases and dysregulation in FSGS. We then performed detailed morphometric studies in our Shroom3 knockdown model of diffuse FPE, followed by *in vitro* studies to understand mechanism, and phenotypic studies in the context of podocyte injury to examine the impact of regulation of glomerular volume on podocyte survival. These studies revealed enhanced activation of AMPK with either Shroom3 or Fyn knockdown in podocytes which accounted for reduced podocyte and glomerular volume. We confirmed that the mechanism of AMPK activation in podocytes in this model is via cytoplasmic re-distribution of the AMPK-activating kinase, LKB1. Since AMPK is a major regulator of cell growth and survival by inhibiting cellular protein synthesis and enhancing autophagy, we postulated that AMPK is a key signaling molecule mediating Vglom regulation and podocyte survival in the context of FPE. Consistent with this, we found that inhibition of AMPK signaling in Shroom3 knockdown mice with podocyte FPE increased Vglom and induced podocytopenia. Furthermore, activation of AMPK in mice following nephron loss-induced hypertrophic injury, restricted Vglom and protected from podocytopenia and progression to FSGS.

## Results

### Glomerular morphometry shows significantly lower Vglom in MCD vs FSGS cases

The Neptune consortium is the largest multicenter prospective cohort of NS collated in the USA with uniform sample/data collection (16). A subset cohort of biopsy proven NS cases currently has available Aperio-scanned images enabling morphometric evaluation by the NEPTUNE pathology core (5). We investigated glomerular volume in this subset with annotated diagnoses of FSGS, MCD or MN (n=80). We excluded “other NS” diagnosis from our analyses due to heterogeneity of this entity. The clinical and demographic characteristics of this cohort at enrollment and the outcomes of these patients by diagnosis are shown in **Table 1**. We applied the Weibel-Gomez formula (see **Fig S1A** (adapted from (5)) and methods) on area cross-sections of glomerular tufts identified in the two paraffin sections (3-60 glomeruli/patient) to perform Vglom estimation. Mean Vglom (μm^3^) calculated from these 2 random sections within the same biopsy were highly correlated validating the morphometric assessment (R^2^=0.987; P<0.001; **Fig S1B**). We then utilized data from all glomerular profiles from one PAS section for mean Vglom estimation and analyses. We identified that MCD cases (n=27) had significantly lower mean-Vglom than FSGS (n=38) or Membranous Nephropathy (MN; n=15) [**Fig 1**A]. To minimize confounding of mean Vglom by older age in FSGS cases, we restricted our analyses to pediatric cases alone (≤18 years). Consistently, pediatric MCD (n=18) also had lower mean Vglom than pediatric FSGS (n=13) [**Fig 1B**]. There were no pediatric MN cases. Furthermore, in Cox proportional hazard models agnostic to diagnoses, increasing Vglom associated with increased composite risk of ESRD and/or 40% or greater decline in eGFR during follow-up, independent of age (HR=1.18 per 10^6^ μm^3^; **Table 2**). The covariates included in these models were those identified as significantly different between the three NS diagnosis in univariable analyses (**Table 1)**. Hence, these data identify lower Vglom in MCD cases vs FSGS representing maintained Vglom regulation in MCD (and dysregulation in FSGS) and, show the association of higher Vglom with adverse renal outcomes in NS.

**Table 1.** Clinical and Demographic data of patients with glomerular morphometry from NEPTUNE consortium.

| Variable | MCD<br>n=27<br>(mean±SD) | FSGS<br>n=38<br>(mean±SD) | MN<br>n=15<br>(mean±SD) | *p-value |
| --- | --- | --- | --- | --- |
| Age at biopsy | 19.11±18.02 | 34.89±23.09 | 52.67±12.92 | < **0.001** |
| Pediatric/Adult | 17/10 | 12/26 | 0/15 | < **0.001** |
| Gender (F/M) | 12/15 | 7/31 | 8/7 | **0.019** |
| UPCR at diagnosis | 4.64±5.96 | 4.38±6.02 | 7.64±6.04 | 0.192 |
| eGFR at diagnosis | 103.58±27.88 | 67.48±31.70 | 72.47±29.96 | < **0.001** |
| **Ancestry (Race)** |  |  |  |  |
| Caucasian | 11 | 17 | 11 | 0.225 |
| African American | 13 | 14 | 3 |  |
| All others | 3 | 7 | 1 |  |
| Immunosuppression diagnosis (Yes/No) | 20/7 | 7/31 | 3/12 | < **0.001** |
| Follow up (months) | 51.22±12.04 | 50.95±8.16 | 50.13±9.05 | 0.941 |
| **Outcomes** |  |  |  |  |
| eGFR slope (per year) | -2.97±6.17 | -3.43±5.13 | -5.45±7.38 | 0.418 |
| Death | 1 | 2 | 0 | 0.662 |
| ESRD | 0 | 6 | 1 | 0.081 |
| ESRD or eGFR decline>40% | 4 | 12 | 5 | 0.250 |

Bx=Biopsy; F= female, M= Male; eGFR = estimated glomerular filtration rate by Modification of diet for Renal disease equation, UPCR = Urine Protein:Creatinine ratio (gram/gram)
\*=p value for Categorical variables obtained by Chi-square and for Continuous variable by ANOVA or Kruskal Wallis

**Figure 1.**
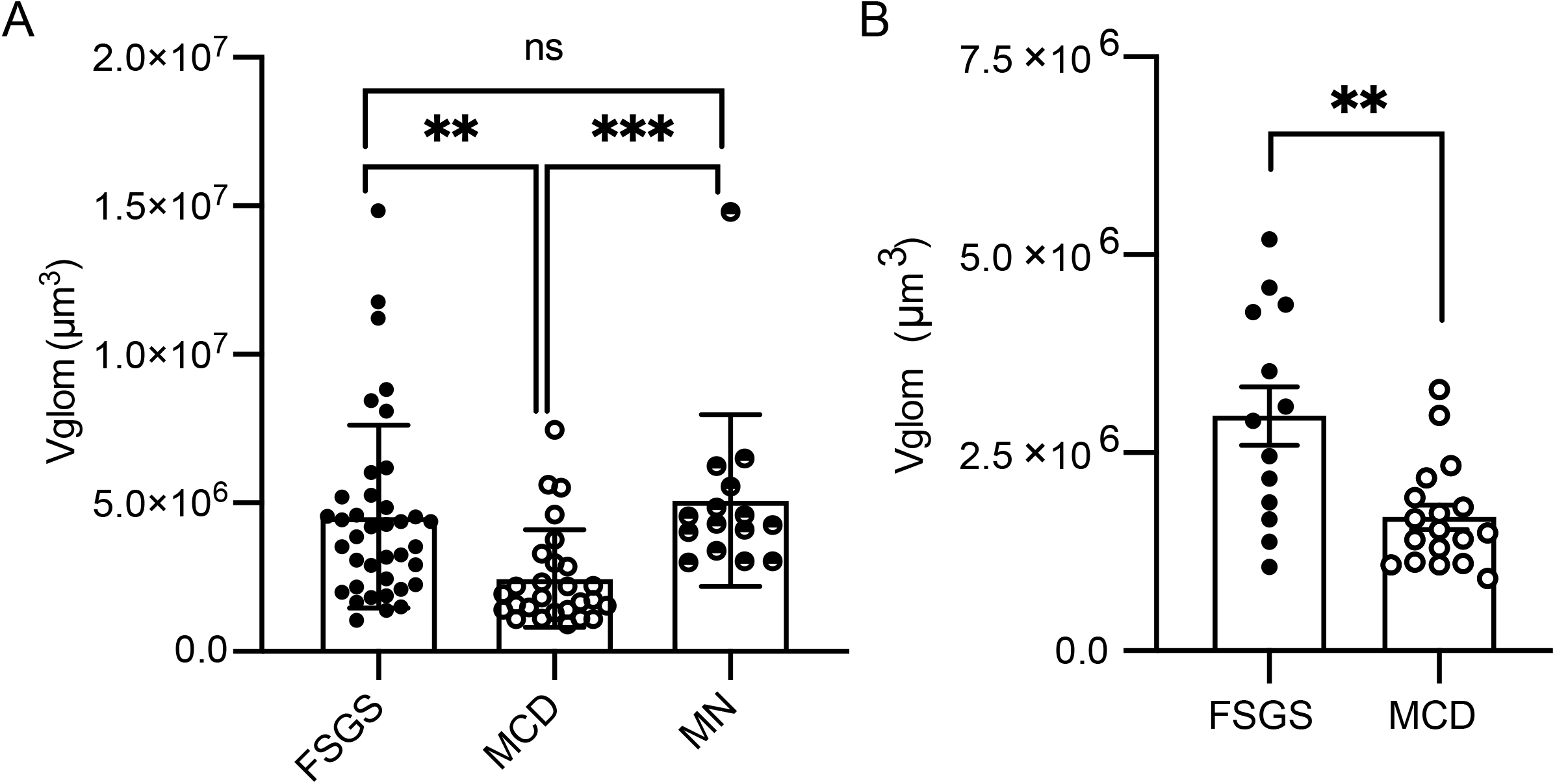
Glomerular morphometry shows significantly lower Vglom in MCD vs FSGS cases. Glomerular morphometry was performed using glomerular area profiles and Weibel-Gomez equation on scanned images of PAS-stained paraffin sections from NS biopsies at enrollment in a subset of the NEPTUNE cohort (n=80 biopsies; 27 MCD, 38 FSGS, 15 MN. Dot plots compare (A) mean Vglom (μm^3^) all FSGS (black solid circles), MCD (black hollow circles) and MN cases (black partly solid circles) in the NEPTUNE morphometry cohort and (B) mean Vglom (μm^3^) in pediatric cases (≤18 years) of FSGS (n=13) and MCD (n=18). [Line/Whiskers=Mean/SEM; Kruskal Wallis test with post test Tukey (>2 groups) Mann-Whitney U test (2 groups); *= P<0.05, **= P<0.01, ***= P<0.001; PAS=periodic acid Schiff].

**Table 2.** Adjusted Cox proportional hazard model for association with renal outcomes.

| Covariate | Model 1 |  |  |  | Model 2 |  |  |  |
| --- | --- | --- | --- | --- | --- | --- | --- | --- |
|  | HR | 95% CI |  | P-value | HR | 95% CI |  | P-value |
|  |  | Lower bound | Upper bound |  |  | Lower bound | Upper bound |  |
| Vglom (per $10^6 \mu\text{m}^3$) | 1.185 | 1.010 | 1.391 | **0.037** | 1.165 | 0.993 | 1.366 | 0.062 |
| Adult (Reference: Pediatric) | 3.693 | 0.596 | 22.870 | 0.160 | 0.319 | 0.051 | 1.975 | 0.219 |
| Age (years) | 1.034 | 1.000 | 1.070 | 0.052 | 1.027 | 0.990 | 1.065 | 0.151 |
| Gender (Reference: Male) | 0.504 | 0.171 | 1.483 | 0.213 | 0.557 | 0.183 | 1.695 | 0.303 |
| Immunosuppression at diagnosis (Reference: Yes) | 2.382 | 0.699 | 8.110 | 0.165 | 2.896 | 0.773 | 10.848 | 0.114 |
| eGFR at diagnosis (ml/min) | NA | NA | NA | NA | 0.990 | 0.971 | 1.009 | 0.311 |

CI= Confidence interval; Vglom= Glomerular volume obtained from area-profile method and Weibel-Gomez formula; eGFR = estimated glomerular filtration rate.

### Global or Podocyte specific Shroom3 knockdown reduced glomerular and podocyte volume

Our human data provided rationale to comprehensively evaluate glomerular morphometry in our inducible *Shroom3* knockdown mice, where we had observed diffuse FPE without podocytopenia, along with reduced podocyte volume. In this mouse model (called “Shroom3-KD”), universal small hairpin RNA (shRNA)-mediated Shroom3 knockdown and turbo green fluorescent protein (tGFP) production, were induced in all tissues by Doxycycline feeding (DOX; see supplemental methods) (10, 17, 18). Non-transgenic or mono-transgenic littermates were used as controls. As we previously reported, global Shroom3-KD mice develop diffuse podocyte FPE by 6 weeks of DOX (10). We first evaluated Vglom using the Cavalieri principle **(Fig S2A-B**) after inducing *Shroom3* knockdown. Here, we identified significant Vglom reductions in Shroom3-KD vs controls **(Fig 2A & Table S1A**; n=8; mean difference in Vglom ∼23%). As described previously (10, 19), we also estimated volume of glomerular components i.e., podocytes (PodoVglom), capillary lumens + endothelium (Cap+EndoVglom), and mesangial components (see methods and **Fig S2C-D**). Glomerular component volume analysis revealed reductions in PodoVglom (p<0.05), and Cap+EndVglom (p=0.06) in Shroom3-KD glomeruli, vs controls **(Table S1A)**. No podocytopenia was identifiable at 6-wks DOX in Shroom3-KD **(Fig 2B)**; indeed, podocyte numerical density (podocytes per unit glomerular volume expressed as n/µm^3^) was higher in Shroom3-KD glomeruli (**Fig S2E**). As previously described, Shroom3 knockdown induced increased albuminuria **(Fig 2C)** without azotemia **(Fig S2F)**. We also observed significantly lower single kidney weights in Shroom3-KD mice, while body weights remained similar to controls (**Fig 2D and S2G**, respectively). The mean difference in kidney weights was 24%, similar to Vglom changes, and suggested the involvement of non-glomerular kidney cells in the renal phenotype observed with global Shroom3 knockdown.

**Figure 2.**
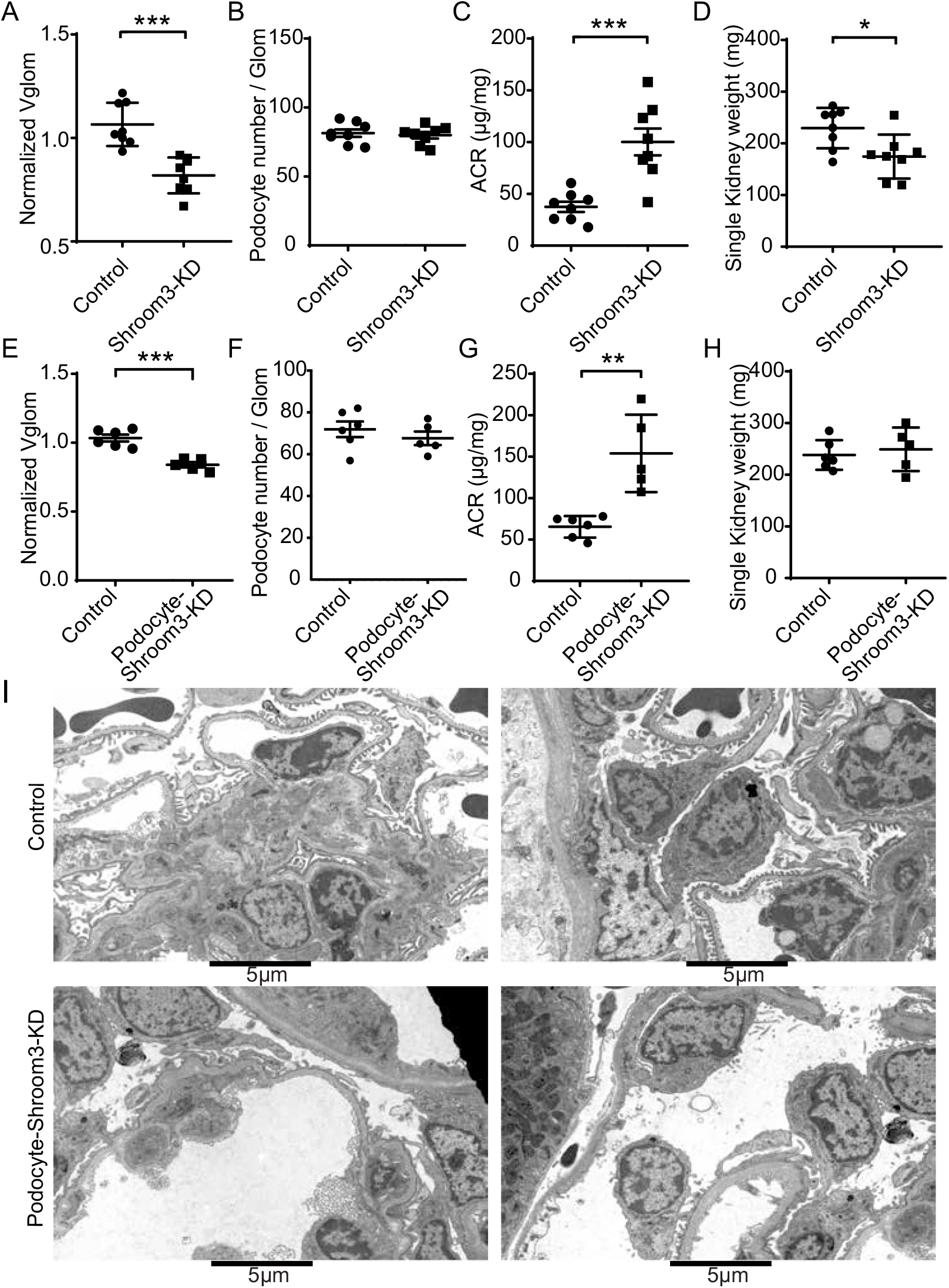
Global or Podocyte specific Shroom3 knockdown reduced glomerular and podocyte volume. (**A-D**) Global Shroom3-KD and control mice (∼12 weeks) were DOX fed for 6-wks (n=8 each). Kidney Tissues were embedded in plastic and 1- μm sections stained with Toluidine blue. Dot-plots compare (A) mean Vglom (X1000 μm^3^) by Cavalieri method (10 Glomeruli/mouse), (B) mean podocyte numbers per glomerulus per animal (fractionator-disector method), (C) Albumin:Creatinine ratio (mcg/mg) at 6 weeks, and (D) Single kidney weights at sacrifice (mg), respectively, in knockdown vs control groups. **(E-I)** To induce podocyte-specific Shroom3 knockdown mice and controls (∼12 weeks age) were DOX fed for 6-wks (n=6 vs 5). Dot-plots compare (E) mean Vglom (X1000 μm^3^) by Cavalieri method (10 Glomeruli/mouse), (F) mean podocyte numbers per glomerulus per animal (fractionator-disector method), (G) Albumin:Creatinine ratio (mcg/mg) at 6 weeks and, (H) Single kidney weights at sacrifice (mg) respectively in Podocyte specific knockdown vs control groups. (I) Panel shows electron microscopic images of representative Podocyte specific Shroom3 knockdown mice and controls (n=2 each) showing podocyte FPE in knockdown glomeruli. [Line/Whiskers=Mean/SEM; Unpaired T-test*= P<0.05, **= P<0.01, ***= P<0.001].

To test the hypothesis that podocyte Shroom3 regulated glomerular volume, we crossed Nphs1-rtTA (20) mice with our inducible Shroom3-shRNA mice (10, 17) for podocyte-specific *Shroom3* knockdown (called “Podocyte-Shroom3-KD” mice). Glomerular protein extracts confirmed Shroom3 knockdown and tGFP production in Podocyte-Shroom3-KD mice **(Fig S2H)**. Podocyte-Shroom3-KD mice with 6-weeks DOX, similarly demonstrated significantly reduced mean Vglom (mean difference ∼15%) as well as reduced PodoVglom and Cap+EndoVglom (**Fig 2E** and **Table S1B)** vs controls (n=6 vs 5). No podocytopenia was identifiable by fractionator-disector method **(Fig 2F)**. Podocyte Shroom3-KD mice also showed increased albuminuria without azotemia (**Fig 2G** and **S2I)**. Neither body weights **(Fig S2J)** nor single kidney weights **(Fig 2H)** were significantly different between Podocyte-Shroom3 KD and control mice, suggesting minimal effect on non-glomerular cells due to podocyte-specific Shroom3 knockdown. EM examination revealed podocyte FPE **(Fig 2I)**, similar to global Shroom3-KD animals (10). These data suggested that, in addition to inducing albuminuria with FPE, global or podocyte-specific Shroom3 knockdown reduced glomerular volume in adult mice without podocytopenia.

### Shroom3 knockdown restricted glomerular hypertrophy post-Unilateral nephrectomy

Since morphometric data from podocyte-specific Shroom3-KD phenocopied global Shroom3-KD, we used global Shroom3-KD mice, which were backcrossed into susceptible BALBc background, for further experiments. To further examine Vglom regulation by Shroom3, we performed unilateral nephrectomy in Shroom3-KD and control mice as described (8), and evaluated glomerular hypertrophy using nephrectomized- and remnant-kidneys (**Fig 3A)** at one-week post-nephrectomy (n= 4 vs 5, respectively). First, after unilateral nephrectomy, mean weight of the remnant-kidney was reduced in global Shroom3-KD vs control animals **(Fig 3B)**. By morphometry, percent change in mean Vglom after nephrectomy was restricted in Shroom3-KD but not in control mice (**Fig 3C)**. As described previously (**Fig S3A-D**) (10, 19), nephrectomized- and remnant-kidneys were used to evaluate the post-nephrectomy expansion of glomerular components. Among glomerular components, PodoVglom expansion and Cap+EndoVglom expansion were significantly restricted (**Fig 3D**) in Shroom3-KD remnant-kidneys vs controls. No podocytopenia was identifiable in remnant-kidneys in either Shroom3-KD or control groups at 1-week post-nephrectomy (fractionator/disector method **(Fig 3E)** or WT1 immunofluorescence (data not shown)). Post-nephrectomy, Shroom3-KD mice had significantly increased albuminuria, but no azotemia vs controls (**Fig S3A-B)**. These data show that remnant-kidneys in Shroom3-KD showed restricted Vglom expansion without podocytopenia, and further demonstrate the regulation of glomerular volume by Shroom3 knockdown.

**Figure 3.**
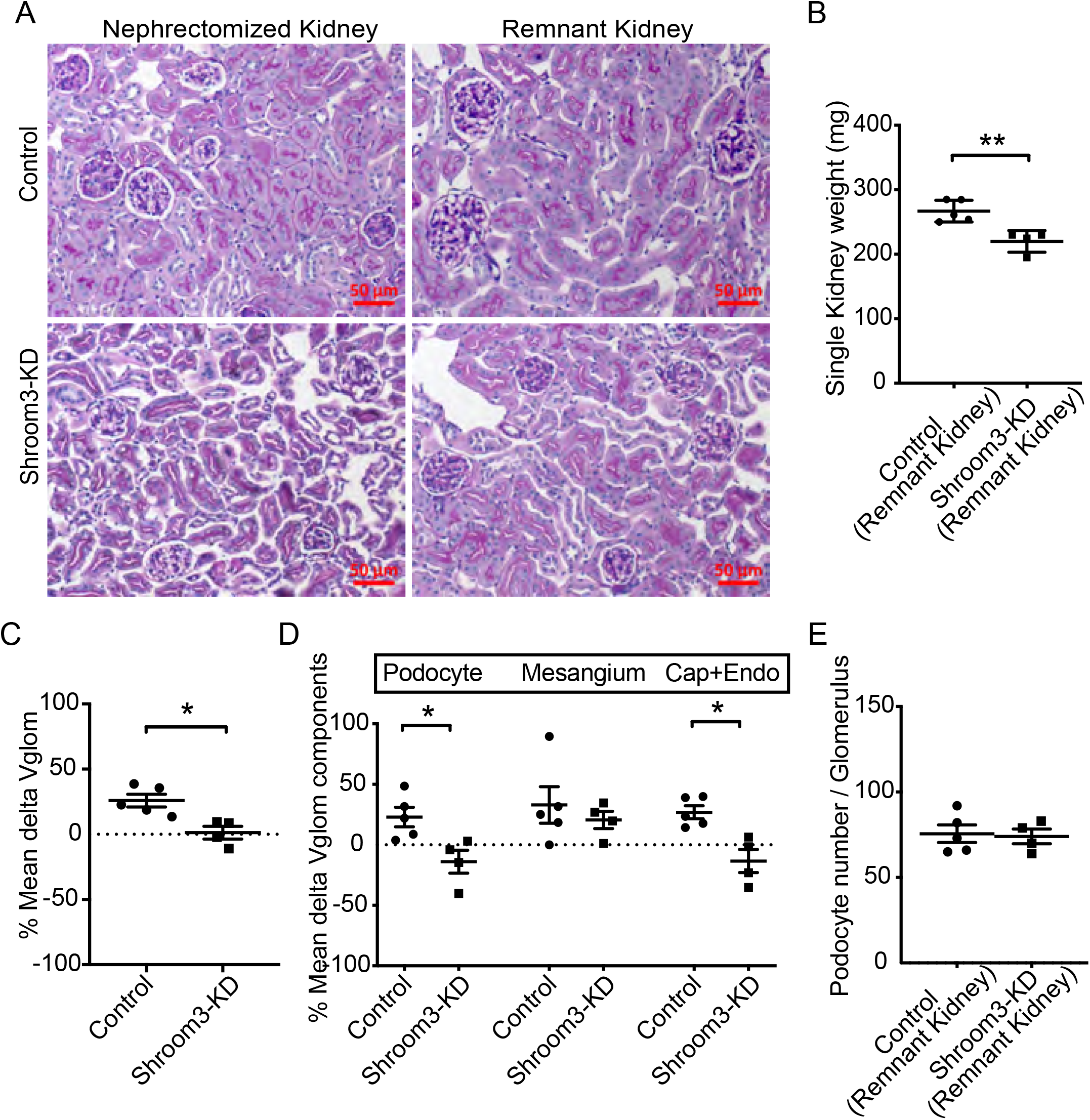
Shroom3 knockdown restricted glomerular hypertrophy post-unilateral nephrectomy. Global Shroom3-KD and control mice (∼12 weeks) were DOX fed for 6-wks and then subjected to unilateral nephrectomy (n=5 vs 4). (A) Panel shows respective representative PAS stained images (20X) of nephrectomized and remnant-kidneys (at day 7 post nephrectomy), of knockdown and control mice. (B) Dot plots compare mean kidney weight of the Remnant kidney (mg) in knockdown vs control groups. (C) Percent change (or delta) of mean Vglom and (D) Vglom components (podocyte, mesangial and capillary + endothelium (Cap+Endo) components), of nephrectomized and remnant kidneys for each animal are shown in dot plots. (E) Dot plots show mean podocyte numbers per glomerulus of remnant kidney per animal in each group (fractionator-disector method, 10 glomeruli/animal) [Line/Whiskers=Mean/SEM; unpaired t-test*= P<0.05, **= P<0.01].

### Shroom3 knockdown reduces cellular protein content and RNA biogenesis *in vitro* and *in vivo* mediated via FYN

Since we reported that Shroom3 knockdown reduced podocyte volume, and inactivated FYN in podocytes (10), we first examined whether the regulation of cell volume by Shroom3 was mediated via FYN. We generated a FYN knockdown stable podocyte line using lentivirally transduced shRNA. *FYN*-shRNA podocytes had significantly reduced cell volume by flow-cytometric forward scatter vs corresponding Scramble-2 podocytes, and similar to Shroom3-shRNA podocytes (10) (**Fig 4A-B**; n=3 sets). This suggested that FYN was downstream of the regulation of podocyte volume by Shroom3.

**Figure 4.**
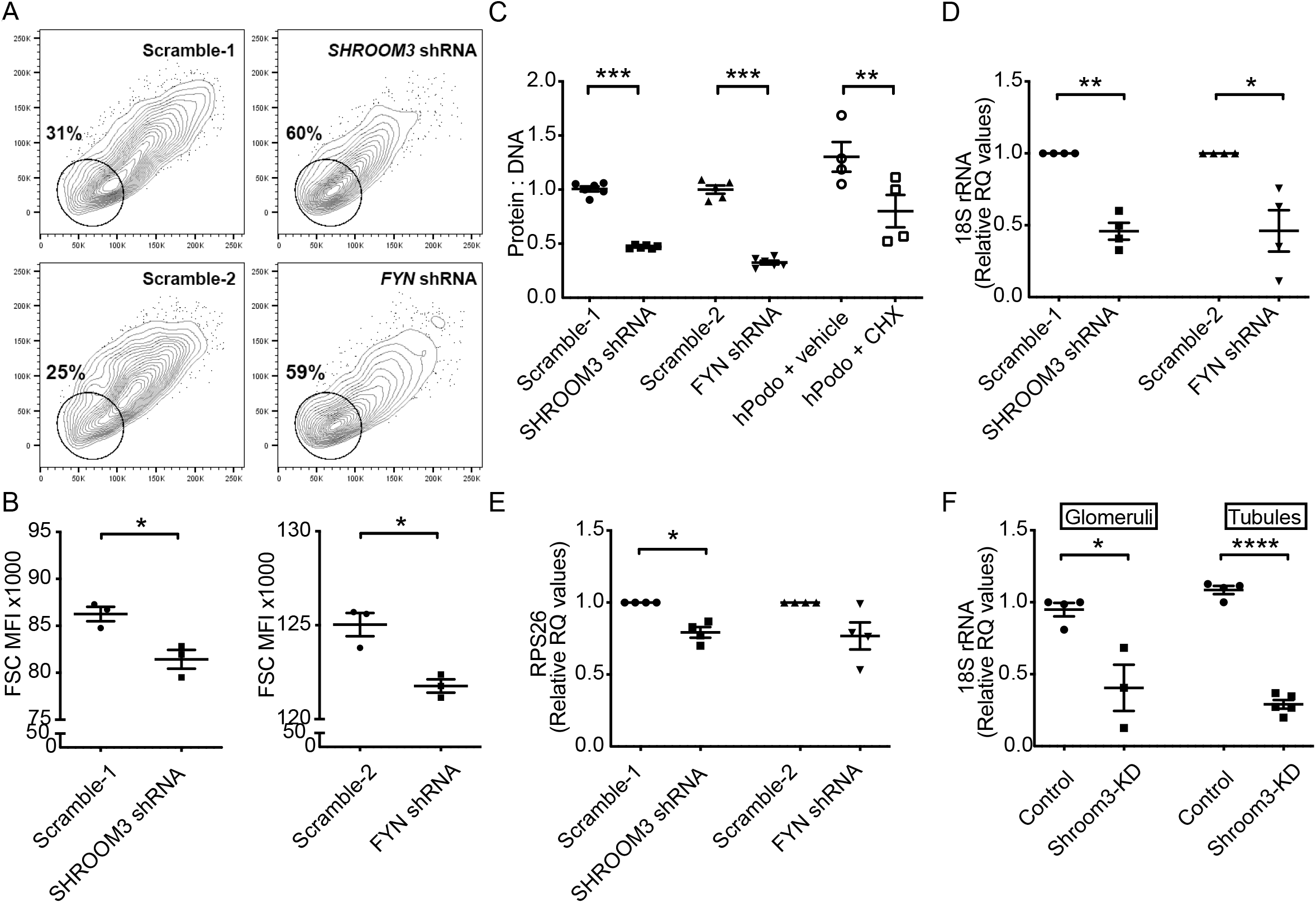
Shroom3 knockdown reduces cellular protein content and RNA biogenesis *in vitro* and *in vivo* mediated via FYN. Puromycin-selectable, stable Shroom3 and Fyn knockdown podocytes were generated using lentiviral shRNA infection (Scramble-1 and Scramble-2 are respective scramble-sequence infected controls). Stable podocytes were differentiated (>7 days) in collagen coated plates. (A) Representative panels show Forward scatter (FSC) MFI of Scramble-1, Scramble-2, Shroom3- and FYN-shRNA podocytes, while (B) shows corresponding dot plots. Arbitrary gate (circle) shows compact clustering of knockdown cells vs scramble (n=3 sets each). (C) Dot plots compare Protein:DNA ratios of Scramble-1, Scramble-2, Shroom3- and FYN-shRNA podocytes. Cycloheximide (CHX) is used as positive control to inhibit protein synthesis. Dot plots show mean fold changes of (D) 18S rRNA copies, and (E) Ribosomal protein S26 (RPS26) transcripts in Scramble-1, Scramble-2, Shroom3- and FYN-shRNA (normalized to Actin; n=4 sets), while (F) compares 18S rRNA copies in glomerular and tubular fractions of control vs Shroom3-KD mice. [Line/Whiskers=Mean/SEM; unpaired t-test *= P<0.05, **= P<0.01, ***= P<0.001; ((D) & (E) compared by paired t-test); RQ=Relative quantity].

To examine whether Shroom3/Fyn knockdown in podocytes reduced cell size by reducing cellular protein content, we performed protein:DNA ratio estimation in Shroom3/Fyn knockdown podocytes vs Scramble-controls. We used Cycloheximide (21), a protein synthesis inhibitor, as a positive control. We identified markedly reduced protein:DNA ratio with Shroom3- or Fyn-knockdown (**Fig 4C**). Next, we examined RNA biogenesis by quantifying ribosomal RNA copies including 18S, 5S and RPS26 in knockdown cells (normalized to Actin). We observed significantly reduced 18SrRNA copies *in vitro* in Shroom3 or FYN knockdown podocytes (**Fig 4D**). *RPS26* transcripts were also reduced in SHROOM3-shRNA podocytes **(Fig 4E)**. *In vivo* in both glomerular and tubular fractions, 18S rRNA copies were significantly reduced in Shroom3-KD kidneys vs Controls (**Fig 4F**) also suggesting inhibited protein synthesis in non-glomerular kidney cells in global Shroom3-KD animals. Interestingly, 5S rRNA transcripts were unchanged *in vitro* Shroom3/Fyn knockdown as well as *in vivo* Shroom3-KD vs Control animals **(Fig S4A-S4B)**.

### Shroom3- or Fyn knockdown increase cellular AMPK activation

Since cellular size and protein biosynthesis were reduced with Shroom3-Fyn knockdown, and prior data from Fyn knockout mice showed increased activation of AMPK (22) - a negative regulator of cellular protein biosynthesis, we examined AMPK signaling after Shroom3 or Fyn-knockdown in podocytes. We identified significantly increased AMPK phosphorylation at Threonine-172 (or pAMPK) in both *SHROOM3* and *FYN*-shRNA transduced podocytes vs respective Scramble controls (**Fig 5A-B**; n=4 sets). Cellular AMPK-activation is stereotypically induced by increased AMP:ATP ratio and is a negative regulator of protein synthesis (23),(24). Consistent with this, phosphorylated-EF2:total EF2 ratio, downstream of AMPK was enhanced in knockdown podocytes vs controls suggesting inhibited protein translation (**Fig 5A-B**). Phosphorylation of MTOR was however not significantly different in Scramble vs knockdown lines. Increased levels of phosphorylated ULK1, and LC3II, (downstream of pAMPK) were also identified in knockdown podocytes (**Fig 5A-B**). We identified increased LC3-positive vacuoles *in SHROOM3*-shRNA podocytes vs control; while Bafilomycin (25) treatment further accentuated LC3-positive vacuoles in *SHROOM3*-shRNA cells confirming significantly increased autophagic flux **(Fig 5C-D)**. In agreement, increased pAMPK staining was seen in glomeruli of Shroom3-KD vs Control mice (n=4 vs 5 mice; **Fig 5E-F; S5A**). Glomerular lysates from Shroom3-KD/Control animals confirmed increased pAmpk and phospho-Ef2 in Shroom3-KD mice (**Fig 5G-H)**. We also examined whole kidney lysates and tubular extracts of Shroom3-KD/Control animals (n=4 each) and confirmed increased Lc3-II in Shroom3-KD mice **(Fig S5B-D)**, suggesting extension of Ampk-activation to non-glomerular cells with global Shroom3 knockdown. Together these data demonstrate increased cellular AMPK activation after Shroom3 or FYN knockdown with reduced protein synthesis and increased autophagy, leading to reduced cellular protein content.

**Figure 5.**
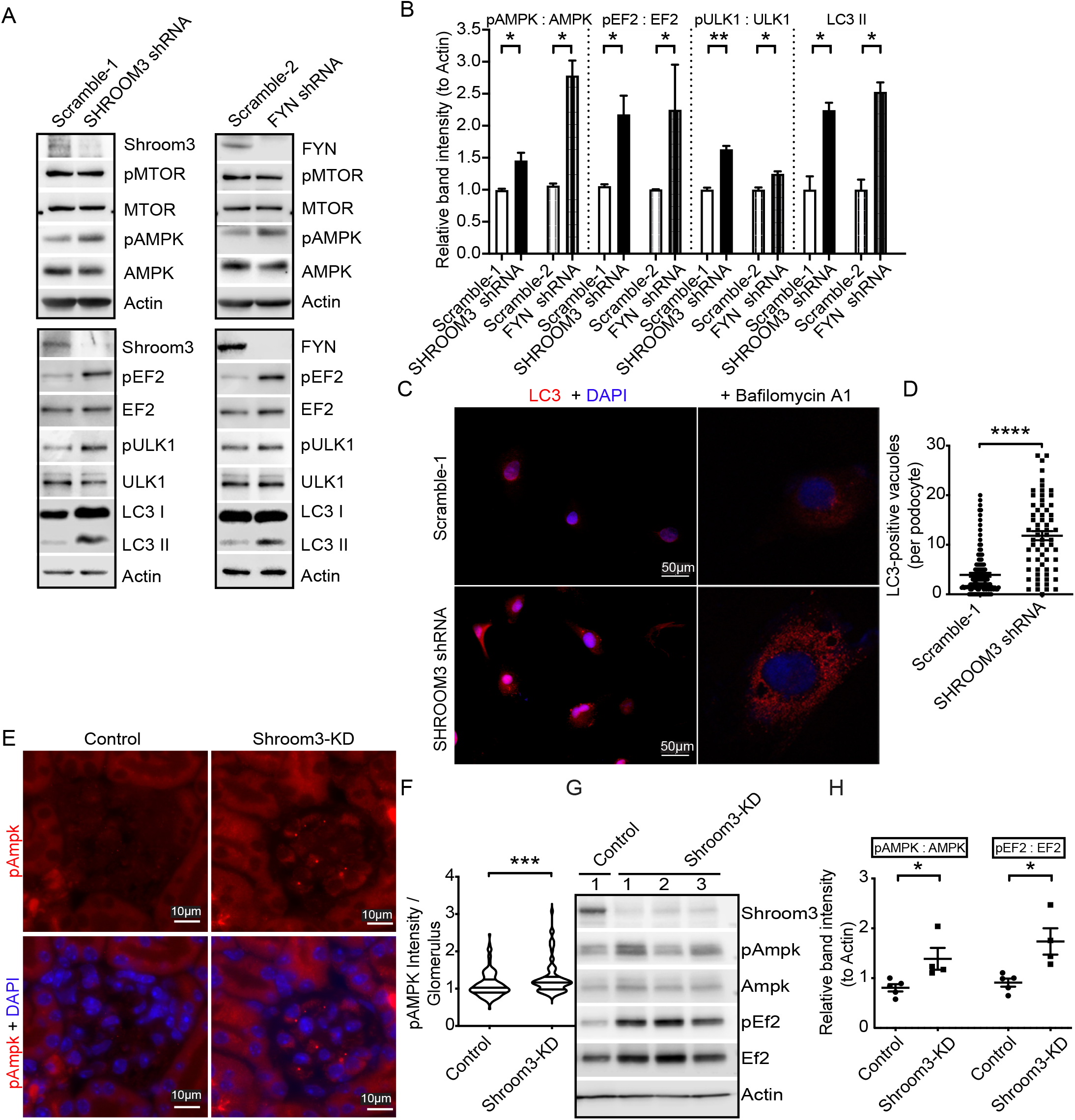
Shroom3- or Fyn knockdown increase cellular AMPK activation. (A) Western blot analysis of lysates from Scramble-1, *SHROOM3*-shRNA (upper panel) and Scramble-2, *FYN*-shRNA podocytes (upper panel) probed for SHROOM3 or FYN and Phosphorylated and Total MTOR, Phosphorylated and Total AMPK, and Actin. Representative WBs of lysates from Scramble-1, SHROOM3-shRNA (lower panel) and Scramble-2, FYN-shRNA podocytes (lower panel) probed for SHROOM3 or FYN, Phosphorylated and Total EF2, Phosphorylated and Total ULK1, LC3-I and II and Actin. (B) Bar graphs show respective relative band intensity (normalized to Actin) (n=4 sets). (C) Representative immunofluorescent images (left panels=20X) of *SHROOM3*-shRNA and Scramble-1 cells stained for LC3 (TRITC). Right panels show confocal images obtained after 24 hours of bafilomycin treatment. (D) Dot plots quantify LC3 positive vacuoles per podocyte (n=2 sets). (E) Representative immunofluorescent images show glomerular staining for phosphorylated AMPK (upper row), and merge with DAPI (lower row) in Shroom3-KD and control mice. (F) Violin-plots quantify intensity of Phosphorylated AMPK/per glomerular outline per group (30 glomeruli/animal; n=5 each). (G) Western blot analysis of glomerular lysates from Shroom3-KD and controls probed for Shroom3, phosphorylated- and total Ampk, phosphorylated- and total Ef2 and Actin, and (H) respective relative band intensities are shown (normalized to actin; n>4 each). [Line/Whiskers=Mean/SEM; unpaired t-test; *= P<0.05, **= P<0.01, ***= P<0.001, ****= P<0.0001].

We previously demonstrated that Shroom3 knockdown led to Fyn-inactivation due to loss of Shroom3-Fyn interaction between the respective SH3-binding- and SH3-domains (10). Additionally, cells with FYN deletion or inactivation showed increased pAMPK, via increased LKB1 cytoplasmic distribution (22). LKB1 phosphorylates Thr-172 of the kinase subunit of AMPK and is ubiquitous. We therefore examined LKB1 localization in *SHROOM3*-shRNA podocytes. Nuclear, cytoplasmic and membrane protein extracts after subcellular fractionation showed consistently reduced nuclear pool of LKB1 and increased LKB1 cytoplasmic:nuclear ratio, suggesting LKB1 redistribution to cytoplasm in *SHROOM3-*shRNA podocytes vs scramble (**Fig S5E-F**). Consistent with AMPK activation, phosphorylated-EF2 was also increased in *SHROOM3*-shRNA podocytes in subcellular fractions **(Fig S5E)**. In summary, following Shroom3 knockdown in podocytes, Fyn inactivation leads to LKB1 redistribution to the cytoplasm and consequent AMPK activation (Summary **Fig S5G)**.

### AMPK activation reduces Vglom and mitigates podocytopenia in aged Shroom3 knockdown mice with podocyte FPE

We have previously reported that aged mice (>1-year) with similar duration of Shroom3 knockdown developed podocyte loss and early FSGS (10), distinct from young Shroom3-KD mice. To understand whether this loss of podocyte protection during aging was associated with reduced Ampk activation in response to Shroom3 knockdown (since age-related decline in AMPK activation is also reported elsewhere (26-28)), we studied aged Control and Shroom3-KD mice. Using aged vs young controls, we demonstrated reduced pAmpk in kidney lysates and in glomeruli of aged controls (**Fig6A, S6A-B**) representing age-related decline of Ampk activation in renal tissues. Further, previously seen enhanced pAmpk in young Shroom3-KD mice, was not observed in aged Shroom3-KD (**Fig 6A vs Fig 5E-H**). Hence, Shroom3 knockdown alone was insufficient to enhance AMPK activation in aged kidneys. At 6 weeks DOX, aged Shroom3-KD developed azotemia and podocytopenia (**Fig 6B** & **6C, respectively; n=5 each)**, in contrast from young Shroom3-KD. Mean Vglom was significantly higher in aged knockdown mice, suggesting an inability to regulate Vglom when Shroom3 knockdown was not associated with AMPK activation (**Fig 6D**). PAS staining also showed mesangial expansion in aged knockdown mice **(Fig 6E)**.

**Figure 6.**
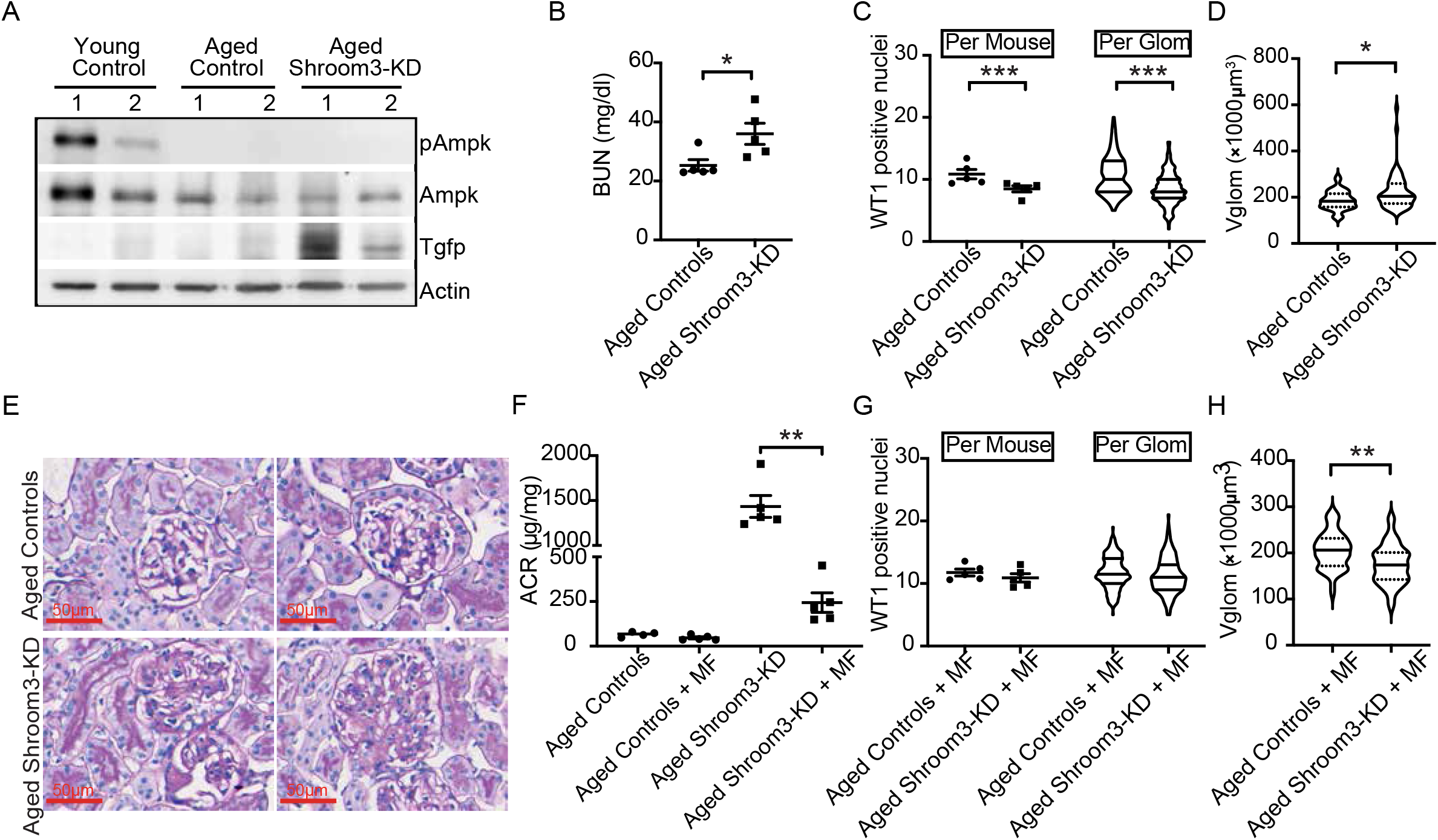
AMPK activation reduces Vglom and mitigates podocytopenia in aged Shroom3 knockdown mice with podocyte FPE. Shroom3-KD and control Mice were aged > 1-year and DOX fed for 6-wks (n=5 per group). (A) Representative western blots of kidney lysates probed for Total and phosphorylated Ampk, Turbo-Gfp, and Actin of young-, aged controls, and aged Shroom3-KD mice. Dot-plots show (B) blood urea nitrogen (mg/dl) in aged control and Shroom3-KD mice. (C) Dot plots show podocytes/ glomerulus/ animal, and Violin-plots (Line at median) show distribution of podocytes/ glomerulus/ group. (D) Vglom per group (x1000μm^3^), between aged control and Shroom3-KD groups. (E) Representative PAS images of two glomeruli (63X) showing mesangial expansion in aged Shroom3-KD mice (lower row) vs aged controls (upper row). In subsequent experiments, Metformin-water (MF) was added at week-2 of DOX to aged control/Shroom3-KD mice. (F) Dot-plots compare Albumin:Creatinine ratio (μg/mg) at 6-wks DOX in aged control vs aged Shroom3-KD mice, both with and without MF treatment. (G) Dot plots compare podocytes/glomerulus/animal and Violin-plots (Line at median) show distribution of podocytes/glomerulus/group and (H) Vglom (x1000μm^3^) per group, between Aged Control+MF and Shroom3-KD+MF groups, respectively [Line/Whiskers=Mean/SEM; unpaired t-test; *= P<0.05, **= P<0.01, ***= P<0.001; PAS=periodic acid Schiff; WT1= Wilm’s Tumor-1 protein] (30 glomerular profiles/animal were used for WT1- immunofluorescence, while 10 glomeruli/ animal-were measured for Vglom).

In subsequent experiments, we used an AMPK activator (29), metformin in drinking water (MF; see methods), to further study the contribution of Ampk to the podocytopenia observed in aged Shroom3-KD (n=5 each group). First, MF treatment restored enhanced Ampk phosphorylation with Shroom3 knockdown in aged Shroom3-KD lysates (significantly greater than in aged controls) (**Fig S6C-D**). Albuminuria in MF-treated aged Shroom3-KD was lower than untreated aged Shroom3-KD (**Fig 6F**) Compared to aged controls, aged Shroom3-KD treated with MF did not show podocytopenia at 6 weeks DOX (**Fig 6G**). MF treatment was also associated with a reduction in mean Vglom in Aged Shroom3-KD vs Aged Controls at 6-weeks **(Fig 6H)**, and thus similar to young Shroom3-KD.

These data suggested that in aged Shroom3-KD mice, loss of Vglom regulation and podocytopenia occurred when podocyte FPE occurred in the absence of enhanced AMPK activation. MF use in aged Shroom3-KD mice enhanced AMPK-activation and reduced Vglom (vs aged controls), improved proteinuria (vs aged knockdown mice), and was protective against podocytopenia.

### AMPK inhibition reverses Vglom reduction and promotes podocytopenia in young Shroom3 knockdown mice

Next, we studied whether pharmacologic AMPK inhibition altered Vglom regulation and reduced podocyte survival in young Shroom3-KD mice with FPE without podocytopenia. We employed Compound C (30) a selective, small molecule, competitive AMPK inhibitor acting via its ATP binding site, and reported to inhibit AMPK activation even in the presence of AMPK activators (31, 32). We administered Compound C at week 5 of DOX feeding (20 mg/kg/dose x 4 doses intraperitoneally) to 8-wk old Shroom3 knockdown mice and controls (n=4 vs 3). We aimed to inhibit AMPK activation after inducing Shroom3 knockdown and podocyte FPE. As shown, Shroom3 knockdown mice had significantly lower body weight after Compound C administration at 8 weeks vs controls (**Fig 7A**). Kidney lysate immunoblotting (**Fig 7B**) and glomerular IF **(Fig S7A-C)** confirmed complete inhibition of Ampk activation by Compound C in both groups of mice. Azotemia was induced by Compound C only in knockdown mice, and not in controls **(Fig 7C vs Fig S2F & S2I)**. Most consistently, morphometry revealed loss of Vglom regulation in Shroom3-KD after Ampk inhibition (**Fig 7D**), with increased Vglom, and Podocyte -, Capillary/ Endothelial-component measurements, vs controls (**Table S2**). Glomeruli of Shroom3-KD mice which previously showed podocyte FPE but without podocytopenia at 6- and 8-wks of DOX (**Fig 1B, 1F** and (10)), now developed podocyte loss (WT1 staining) **(Fig 7E-F)** with Compound C. Hence inhibition of AMPK by Compound C in young Shroom3 knockdown mice, was followed by loss of protective morphometric changes and induction of podocytopenia.

**Figure 7.**
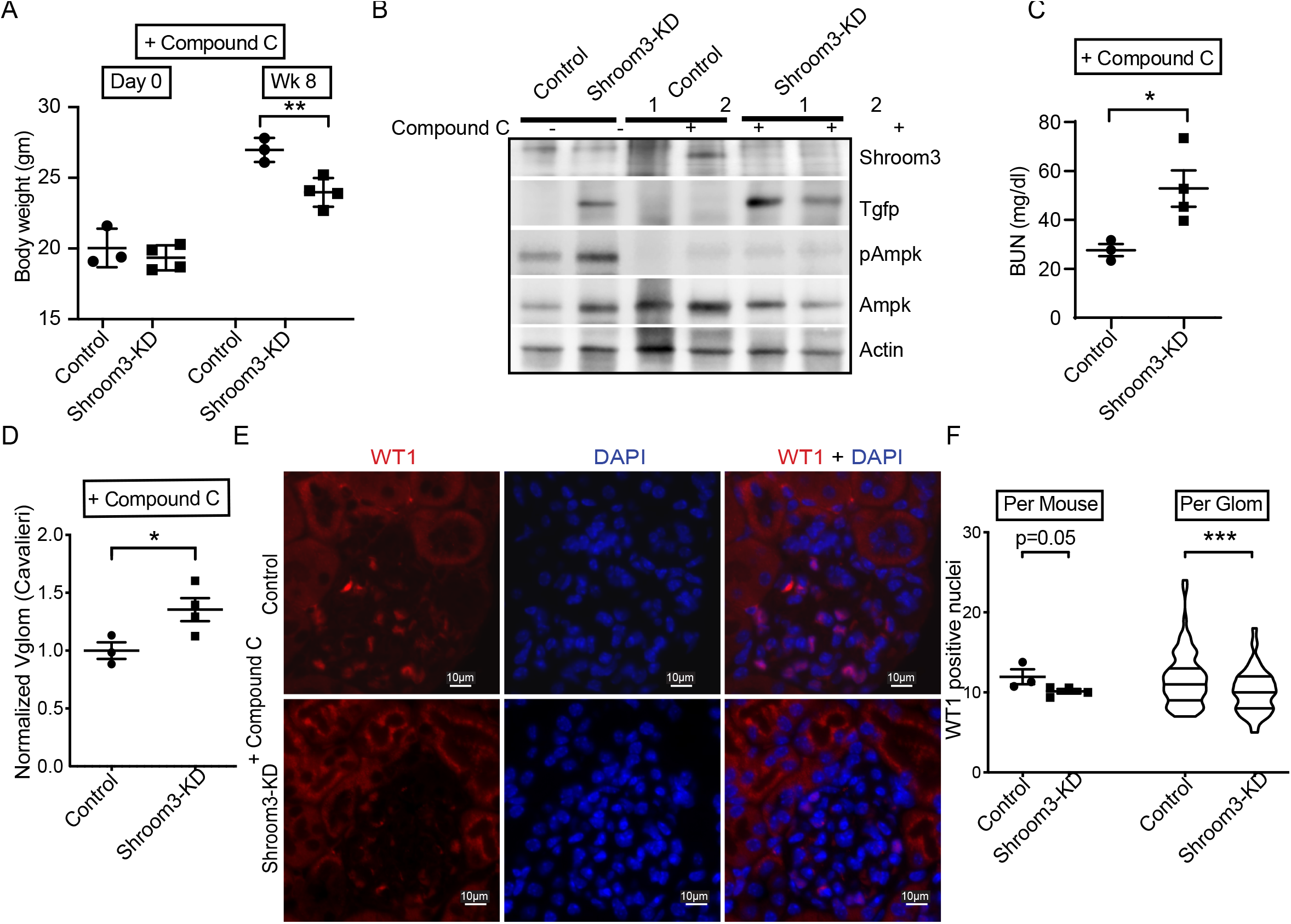
AMPK inhibition reverses Vglom reduction and promotes podocytopenia in Shroom3 knockdown mice. Shroom3-KD and control mice (∼8 weeks old) were DOX fed for 5 wks. AMPK-inhibitor Compound C was injected (20mg/kg intraperitoneally x 4 doses at wk-5) and mice followed till wk-8 when tissues were collected. (A) Dot plots show mean body weight (gms) (day 0 vs at week 8) in both groups. (B) Representative WBs of whole kidney lysates from control/Shroom3-KD mice confirmed inhibition of AMPK activation by Compound C. Dot plots show (C) blood urea nitrogen (BUN in mg/dl) and (D) morphometric quantification of Vglom (x1000μm^3^) in Shroom3-KD and control mice treated with Compound C. (E) Representative immunofluorescent images (40X) of WT1 and DAPI stained glomerulus of Shroom3-KD and control mice receiving Compound C. (F) Dot plots show podocyte numbers/glomerulus/animal and Violin-plots (Line at median) show corresponding distribution of podocytes/ glomerulus/ group (30 glomerular profiles/mouse). [Line/Whiskers = Mean/SEM; unpaired t-test; *= P<0.05, ***= P<0.001; WT1= Wilm’s Tumor-1 protein].

### AMPK-activation reduces glomerular volume and preserves podocyte numbers in nephron loss-induced glomerular hypertrophy

Finally, we asked whether pharmacological Ampk activation would promote favorable Vglom regulation and podocyte survival in wild-type mice. To test this hypothesis, we administered PF0640957 (PF), a highly specific AMPK agonist (33), to BALBc mice subjected to 5/6^th^ nephrectomy - a model for FSGS resulting from maladaptive hypertrophy of remnant glomeruli and podocytes. BALBc mice, without or with (BALBc+PF mice) were subjected to 2/3^rd^ nephrectomy, followed by contralateral nephrectomy seven days later, and sacrificed after a further 6 weeks (n= 6 vs 5, respectively; See methods). PF06409577 gavage was initiated a day before the first surgery.

Baseline BUN was similar in both groups (not shown), and mice showed similar weight loss trends with surgery (**Fig S8A**). Kidney lysates from BALBc+PF group obtained from sequential nephrectomy samples confirmed Ampk activation (**Fig 8A**). BALBc+PF mice showed significantly attenuated albuminuria (**Fig 8B)**, and improved azotemia (**Fig 8C, S8B**) by sacrifice.

**Figure 8.**
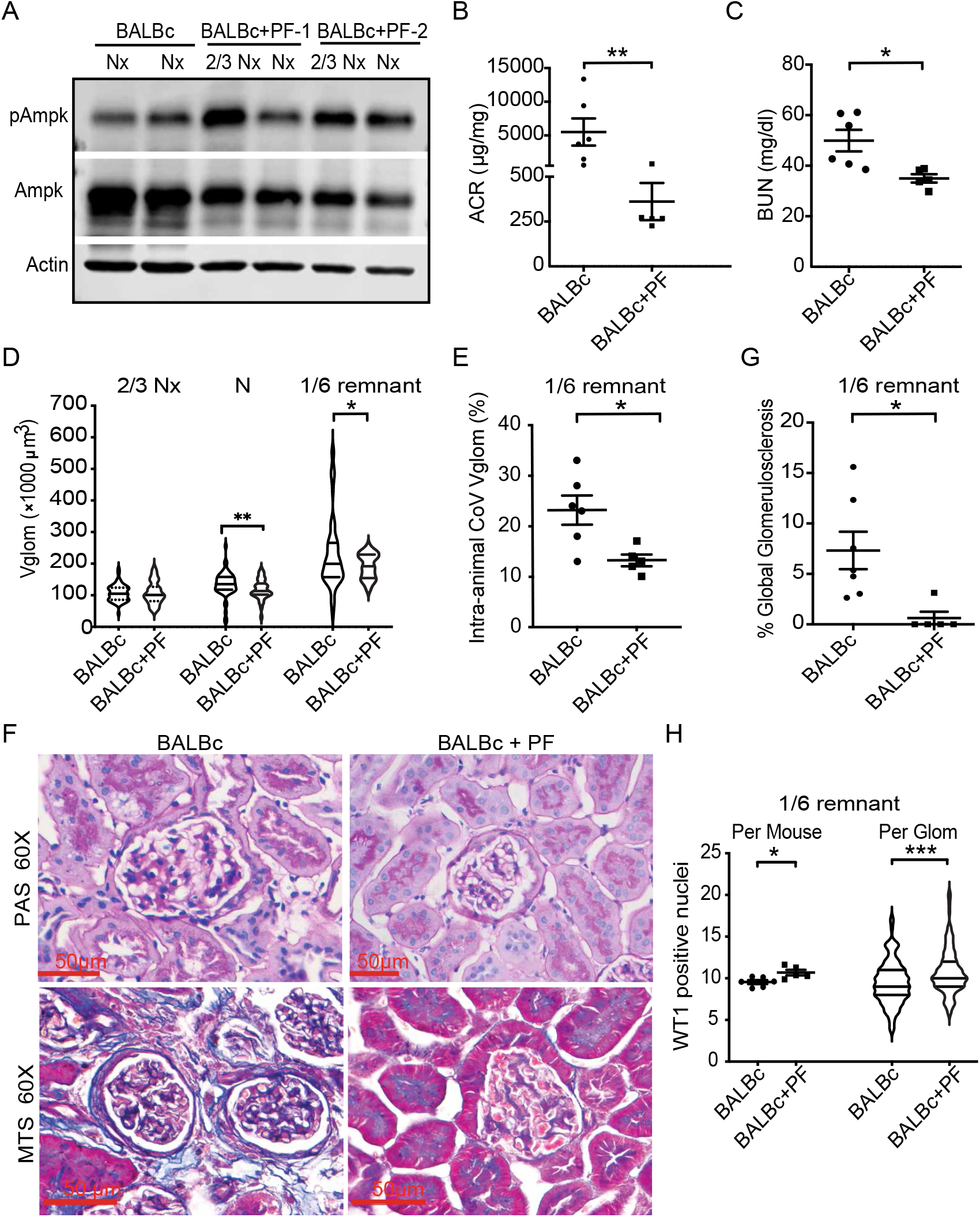
AMPK-activation reduces glomerular volume and preserves podocyte numbers in nephron loss-induced glomerular hypertrophy. Adult BALBc mice (near 8-wks) underwent 2/3^rd^ nephrectomy (2/3 Nx), followed by contralateral nephrectomy (Nx) 1 week later, and were followed for 6 more weeks when the remaining kidney tissue (1/6^th^ remnant) was harvested. Experimental animals were gavaged with AMPK-activator PF06409577 (3 doses/week; BALBc (n=6) vs BALBc+PF (n=5)). (A) Western blot analysis of lysates from 2/3^rd^ Nx and Nx kidneys show AMPK activation in BALBc+ PF vs BALBc. Dot-plots compare (B) Albumin:Creatinine ratio (μg/mg) and (C) blood urea nitrogen (mg/dl) at sacrifice. (D) Violin-plots show the distribution of Vglom (x 1000μm^3^) in the 2/3^rd^ Nx, Nx, and 1/6^th^ remnants of BALBc and BALBc+PF animals (Line at median; n=10 glomeruli/sample). (E) Dot plots show intra-animal coefficients of variation of Vglom in 1/6^th^ remnants, expressed as percentage. (F) Representative images (60X) of 1/6th remnants of PAS- (upper row) and MTS-stained sections (bottom row) show increased glomerulosclerosis in BALBc group. (G) Percentage of glomeruli in 1/6^th^ remnants with global sclerosis (on MTS stain) are shown. (H) Dot plot shows podocytes/ glomerular profile/ animal by WT1/DAPI stain, and Violin-plots (Line at median) show distribution of podocytes/ glomerulus in each group (≥20 glomerular profiles per 1/6^th^ remnant). [Line/Whiskers = Mean/SEM; unpaired t test; *= P<0.05, **= P<0.01, ***= P<0.001; PAS=periodic acid Schiff; MTS=Masson Trichrome; glom=glomerulus; WT1= Wilm’s Tumor-1 protein].

We performed morphometry on the serially obtained 2/3^rd^ kidneys (2/3^rd^ Nx) contralateral nephrectomies (Nx) and 1/6^th^ remnants. Baseline Vglom **(Fig 8D)** and podocyte numbers **(Fig S8C)** from 2/3^rd^ Nx kidneys were similar in both groups. Nephrectomized kidneys as well as 1/6^th^-remnants in BALBc+PF group, showed significantly reduced Vglom vs corresponding BALBc samples **(Fig 8D)**. Further, 1/6^th^ remnants of BALBc showed widely distributed values of Vglom **(Fig 8D)**, suggesting occurrence of both sclerosed and hypertrophic glomeruli, while BALBc+PF remnants demonstrated improved Vglom regulation with significantly reduced intra-animal coefficients of variation **(Fig 8E)**. Histologically, BALBc-remnants had significantly more sclerotic glomeruli on Masson-trichrome stain (**Fig 8F-G)**, whereas BALBc+PF 1/6^th^ remnants showed significantly higher podocyte numbers **(Fig 8H)**, and podocyte numerical density **(Fig S8D)**. Hence, Ampk agonism regulated glomerular volume, restricted Vglom hypertrophy, promoted podocyte survival and preserved kidney function in a volume stress model of glomerular injury in wild-type BALBc mice.

Next, we used another AMPK activator sodium salicylate vs vehicle treated wild-type BALBc controls (n=5 vs 5 mice) in an Adriamycin induced injury model of FSGS. Two vehicle treated animals reached humane endpoints prior to study termination. Further, they exhibited reduced albuminuria, lower creatinine and significantly rescued podocyte numbers as well as markedly mitigated FSGS than controls (**Fig S8E-I**; n=5 vs 3 mice). In summary, we reveal AMPK signaling as a key regulator of Vglom and podocyte survival in injured podocytes with FPE or when facing glomerular hypertrophic stress **(Fig 9)**.

**Figure 9.**
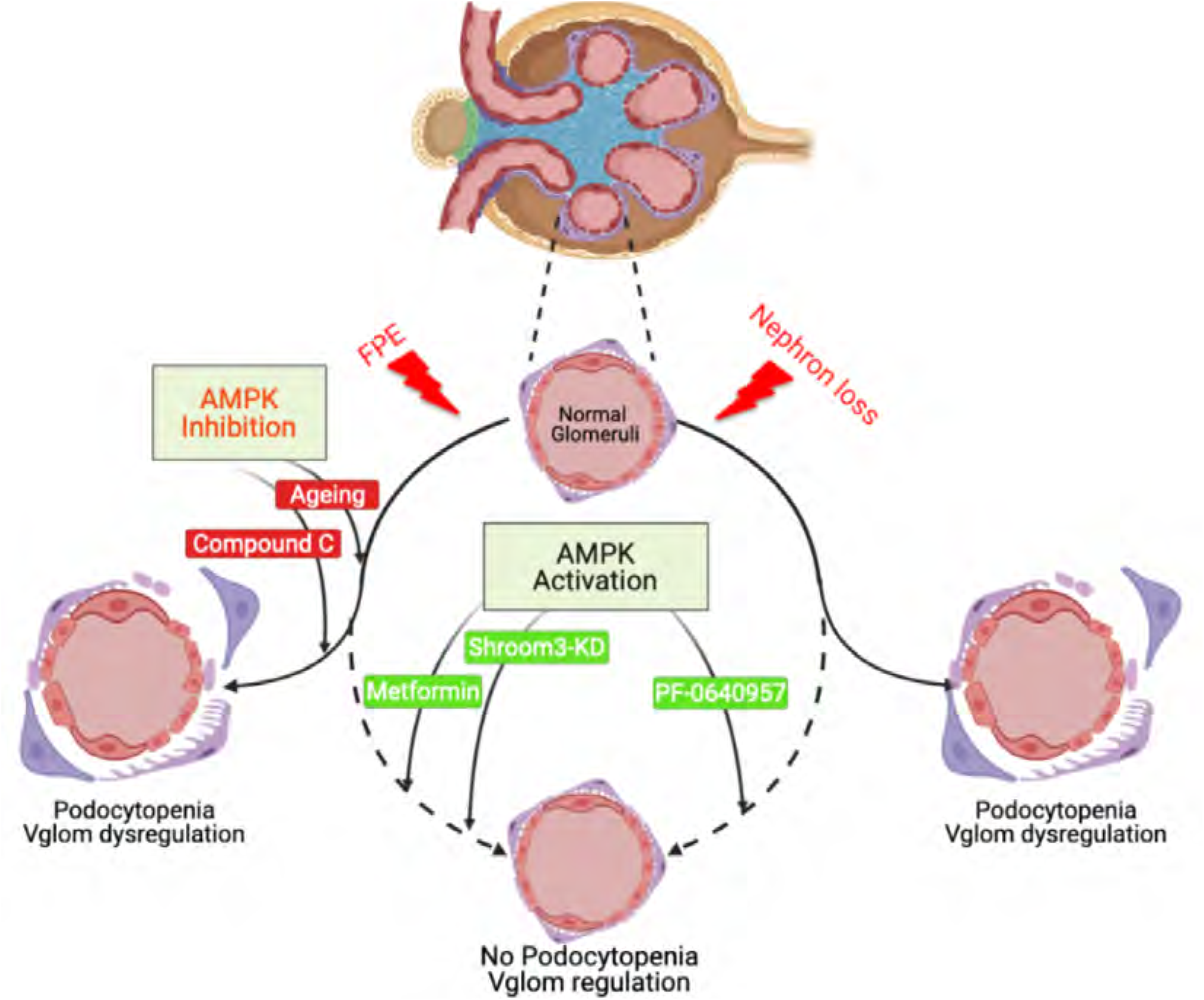
Role of AMPK signaling in glomerular volume regulation and podocyte survival in the context of podocyte injury. Cartoon depicts response of normal glomeruli to injury stimuli i.e., nephron loss in 5/6^th^ nephrectomy or podocyte FPE with Shroom3 knockdown (solid arcuate lines). Sustained hypertrophic stress, or Ampk inhibition (compound C or Ageing shown as red lines) along with podocyte FPE, led to glomerular volume dysregulation and podocytopenia. In these contexts, Ampk activation (green lines) with PF, MF or by Shroom3 knockdown (in young mice), preserved glomerular volume regulation and protected from podocytopenia (arcuate dashed lines), thus promoting an MCD-like pathology in young Shroom3-KD.

## Discussion

Podocyte loss correlates with renal survival in experimental models (8, 34). The association of Vglom and podocyte morphometric parameters with outcomes is reported from human cohorts(3, 5, 35, 36). Hence, identifying novel signals for glomerular volume regulation in injured glomeruli and podocyte survival mechanisms in the presence of FPE, are both crucial. Using a multicenter NS cohort, we first show that MCD diagnosis, where diffuse podocyte FPE does not lead to podocytopenia, is associated with lower Vglom vs FSGS and MN. Furthermore, reduced Vglom by itself was associated with improved renal outcomes. Based on these observations, and analogous findings in our young Shroom3-KD mice with diffuse FPE without podocytopenia, we investigated downstream signaling involved in Vglom regulation and podocyte survival. We identify enhanced AMPK phosphorylation with Shroom3- or Fyn-knockdown via LKB1 release (22). Ampk activation was associated with increased autophagy and reduced cellular protein content downstream. In this regard, we make the important observation here that podocyte specific *Shroom3* knockdown regulates not only PodoVglom, but also Cap+EndoVglom and total Vglom. These data suggest that primary alterations in podocytes without evidence of podocyte loss can regulate adjacent capillary volume. Recent elegant data using mice with hyperactive Mtor in podocytes, similarly showed increases in podocyte size, PodoVglom, as well as total Vglom before the onset of podocyte loss (4). These data relay the need to study the potential crosstalk mechanisms initiated in the podocyte and involved in Vglom regulation.

Ampk activation regulates several downstream signaling pathways including anabolism, autophagy, energy conservation and mitochondrial homeostasis (reviewed by Carling (37)). Among these, identifying the central pro-survival mechanism/s once podocyte AMPK is activated, needs further examination. AMPK regulates cell growth, by modulating mTORC1 signaling, but also by directly inhibiting ribosomal biogenesis, protein translation (by phosphorylating EF2), and activating autophagy (23) (24). We show here that AMPK activation in podocytes regulated podocyte volume, albeit without significant changes in MTOR phosphorylation (4). This is consistent with recent work showing the *in vivo* role of Ampk in the regulation of podocyte autophagy (38, 39). This is also relevant as AMPK-autophagy axis is reported to be protective in FSGS models (38, 40). Reduced 18S RNA seen here with increased pAMPK is likely from inhibition of RNA-polymerase-I by PRKAG2 (gamma-2 subunit of AMPK) also expressed by podocytes (24).

AMPK activation was beneficial to podocyte survival in diabetic and obesity injury models (41, 42). We, and others (43, 44), have now demonstrated that Ampk activation can mitigate FSGS and improve podocyte survival in animal models of FSGS. Our experiments further reveal that in Shroom3-KD with podocyte FPE (10), coincidental AMPK activation regulated glomerular volume and prevented podocytopenia, promoting an MCD-like pathology. The critical role of Ampk activation in preventing glomerular enlargement, podocytopenia and azotemia was demonstrated in both aged mice (in which Ampk activation is reduced) and following pharmacologic Ampk inhibition by Compound C. Most importantly, activating Ampk in podocytes with Shroom3 knockdown induced FPE, or after nephron loss in wild type mice, promoted adaptive Vglom regulation and prevented podocytopenia. While we show morphometric data from MCD, future work is needed to specifically study the role of Ampk signaling in MCD and, in regulating podocyte survival to mitigate progressive NS. Our data are therefore an important platform to examine AMPK based therapeutics in human NS using approved AMPK-activators (45) or novel LKB1-based agents (46).

We have not addressed Fyn knockout *in vivo* (47). We acknowledge this as a limitation of our work. However, global Fyn knockout mice had FPE without podocyte loss at least till 1-year age (similar to Shroom3 knockdown mice) and reduction in cell size, suggesting *in vivo* Ampk activation (13, 22). We cannot completely rule out ancillary mechanisms of Ampk activation by cytoskeletal or metabolic stressors in Shroom3 knockdown cells. Since we used global Shroom3-KD mice, our current data also cannot exclude beneficial effects of Ampk activation (by MF, Salicylate or PF) and deleterious effects of Ampk inhibition in non-podocyte cells as contributing to observed renal phenotypes.

In summary, utilizing a Shroom3 knockdown murine model, we reveal the role of Ampk signaling in the regulation of podocyte and glomerular volume. We applied an ageing model and pharmacologic agents to demonstrate the key role of AMPK in regulating podocyte survival in injured glomeruli with podocyte FPE. These findings are of importance to podocyte biology and pathology in NS and have considerable application to therapeutics in glomerular diseases due to the availability of AMPK activators.

### Concise Methods: (See Supplemental methods)

#### Cell Culture

Human podocyte cell line, were differentiated using RPMI-1640. Protein:DNA ratio assay was done using 1000-podocytes/well in 96 well plates. ***in vitro* autophagy studies** were performed using Bafilomycin A1 at Day 7 of differentiation (100nM for 24-hrs) (25).

#### Reverse transcription/Quantitative-PCR

*Transcript* expression was assayed by real-time polymerase chain reaction using specific primers (see Table S3). Amplification curves were analyzed via the delta-delta CT method.

#### Western Blotting

Cells were lysed for immunoblot analysis as described (19) and probed using antibodies against SHROOM3, FYN, Actin, phospho-mTOR (Ser2448), mTOR (Ser2448), phospho-AMPK (Thr172), AMPK, phospho-eEF2 (Thr56), eEF2, turboGFP, phospho-ULK1 (Ser555), ULK1 (D8H5), LC3A/B.

#### shRNA suppression studies

Human *SHROOM3*(10),(17) and *FYN* short hairpin clones (Dharmacon, inc, USA) were tested for optimal suppression in 293-T cells, HK-2 cells and human podocyes, to generate stable cell lines.

#### Immunofluorescence

5-μm paraffin sections of formalin-fixed kidney-tissues were deparaffinized and processed for unmasking and antigen retrieval followed by incubation with primary antibodies.

#### Quantitative Image Analysis

pAMPK: Glomeruli were outlined using Zen pro (Zen 2.6 (blue edition)) software (40X images), and area of pAMPK-staining were measured for signal intensity and expressed as average signal intensity total area/glomerulus.

#### Murine Shroom3 knockdown model

(Supplemental methods). Double transgenic mice, Chicken beta-Actin promoter, reverse Tetracycline Transactivator (rtTA) /Shroom3-shRNA mice CAGS-rtTA or Nphs1-rtTA (generous gift from Dr Miner to Dr He (20)) were generated for global- or podocyte-specific knockdown (i.e., Shroom3-KD or Podocyte-Shroom3-KD, respectively). Male mice (∼8-12-week-old) were DOX-fed for 6 weeks. For *Ageing studies*, Shroom3-KD and BALBc control mice were aged >1 year, before DOX. Kidney tissues and Glomeruli were processed (Supplemental methods). *Uninephrectomy*: DOX-fed male Shroom3-KD/Podocyte-Shroom3-KD vs control mice were nephrectomized and kidneys saved. Residual kidney (Rx) was collected at 1-wk. *5/6 Nephrectomy*: First surgery was performed to remove 2/3^rd^ of the left kidney. After 7 days contralateral nephrectomy was done. At each surgery tissues were saved for studies (48). *AMPK inhibition studies using Compound C*. After 4-wks DOX, mice were injected with Compound C (20mg/kg x 4 doses), sacrificed at 8-wks. *AMPK activation* studies: In 5/6 nephrectomy mice, PF-06409577 was administered (gavage 50 -100 mg/kg; Supplemental methods) (48) *Metformin protocol*: Metformin (dose= 500 mg/500 ml in drinking water (49)) was added at week-2 DOX. *Sodium Salicylate protocol:* Tail vein injections of Adriamycin at 9.8∼10.4mg/kg were performed in 8-wk old BALBc mice to induce FSGS. Control Group was fed with 3% Sucrose solution (n=5/3) and Experimental Group with 250mg/l Sodium Salicylate in 3% Sucrose solution (n=5) and followed for 2 weeks till termination of the experiment.

#### Glomerular Morphometry (see supplemental methods)

*<u>Glomerular Volume- Weibel-Gomez Method:</u>* The Weibel-Gomez method was used to measure glomerular volume (Vglom) in human NS biopsies. This method uses one PAS-stained and one Trichrome stained paraffin section from Aperio-scanned images of NS biopsies from the NEPTUNE study (5). The areas of all complete glomerular profiles present in the section were measured by planimetry (Figure S1A). Vglom = A^3/2^ x 1.38 µm^3^ where A is the average glomerular tuft area and 1.38 is the shape correction factor assuming glomeruli are spheres (50). The mean-Vgloms obtained from two sections within each patient were highly correlated (Fig S1B). The PAS-stained sections were used for stereological analyses. *<u>Glomerular Volume-Cavalieri Method</u>*: One-millimeter cubes were cut from the cortex, fixed in glutaraldehyde and embedded in epon. Glomerular volume (Vglom) was measured using 1-µm-thick sections and the Cavalieri principle (Fig S2A). As described previously (10, 19), high magnification images (∼1700X) were used for estimation of Vglom components (Supplemental methods; Fig S2B-D), i.e., podocytes (Podo Vglom), capillary lumens + endothelial cells (Cap+Endo Vglom), and mesangium were calculated by measuring the volume fraction of each component and then multiplying each fraction by the glomerular volume. Podocytes were counted using the fractionator/disector method (Supplemental methods (51)). The morphometrist was blinded to disease group for the human biopsies, and to experimental group in murine data.

### Statistical analysis

De-identified clinical information was obtained for the NS morphometry cohort and linked to morphometry using unique-IDs. *For human data*, univariate comparisons of clinical /demographic factors between NS categories were done using ANOVA (Kruskal Wallis for corresponding nonparametric analysis with post-test Dunn’s) for continuous variables, and Chi-Square for proportions. Spearman correlation coefficient was used to compare Vgloms within patient. Cox proportional hazard models were used for multivariable survival associations, including clinic-demographics identified as significantly different in univariable analyses. NEPTUNE determined outcomes of end-stage renal failure, eGFR decline ≥ 40% from baseline, or a composite of these events were evaluated as outcomes. Time from biopsy to event was utilized. For *in vitro and in vivo* experiments, unpaired *t* test (Mann-Whitney test for corresponding non-parametric analysis) was used for data between two groups. Statistical significance was considered with two-tailed *P*<0.05.

### Study approval

Institutional IRB and IACUC approved protocols were available for all human data and mouse experiments performed according to humane endpoints (Supplemental Methods).

## Supporting information

Addendum

## Author contributions

Research/study design: KB, QL, JCH, and MCM. Experimentation: KB, QL, CW, MP, AC, FG, AG. Mouse husbandry: FG, MP, QL, KB, MCM. Morphometry: JMB and KVL. Histology: FS, LL. Analyzing data: KB, QL, WZ, NC and MCM. Providing reagents: AS, LK, BD, BM, JCH. Manuscript: KB, JCH and MCM. All authors contributed to editing of manuscript.

## Acknowledgments

We acknowledge Dr David Carling, University college London, UK, and Drs Michael Caplan, Lloyd Cantley, Shuta Ishibe, Yale University School of Medicine, USA, for critical discussions regarding data, and Professor Peter Heeger, Mount Sinai for feedback on data, Gail Celio, University of Minnesota Imaging Center and Ronald Gordon at the electron microscopy core Icahn School of Medicine, Mount Sinai.

## NEPTUNE Acknowledgements

We acknowledge Tina Mainieri and Jonathan Troost, Dr Holzman and members of the NEPTUNE anciliary study committee, and the NEPTUNE consortium investigators.

## Members of the Nephrotic Syndrome Study Network (NEPTUNE)

NEPTUNE Enrolling Centers

*Cleveland Clinic, Cleveland, OH*: K Dell^*^, J Sedor^**^, M Schachere^#^, J Negrey^#^

*Children’s Hospital, Los Angeles, CA*: K Lemley^*^, B Silesky^#^

*Children’s Mercy Hospital, Kansas City, MO*: T Srivastava^*^, A Garrett^#^

*Cohen Children’s Hospital, New Hyde Park, NY:* C Sethna^*^, K Laurent ^#^

*Columbia University, New York, NY:* P Canetta^*^, A Pradhan^#^

*Emory University, Atlanta, GA:* L Greenbaum^*^, C Wang**, C Kang^#^

*Harbor-University of California Los Angeles Medical Center:* S Adler^*^, J LaPage^#^

*John H. Stroger Jr. Hospital of Cook County, Chicago, IL:* A Athavale^*^, M Itteera

*Johns Hopkins Medicine, Baltimore, MD:* M Atkinson^*^, T Dell^#^

*Mayo Clinic, Rochester, MN:* F Fervenza^*^, M Hogan**, J Lieske^*^, V Chernitskiy^#^

*Montefiore Medical Center, Bronx, NY:* F Kaskel^*^, M Ross^*^, P Flynn^#^

*NIDDK Intramural, Bethesda MD:* J Kopp^*^, J Blake^#^

*New York University Medical Center, New York, NY:* H Trachtman^*^, O Zhdanova**, F Modersitzki^#^, S Vento^#^

*Stanford University, Stanford, CA:* R Lafayette^*^, K Mehta^#^

*Temple University, Philadelphia, PA:* C Gadegbeku^*^, S Quinn-Boyle^#^

*University Health Network Toronto:* M Hladunewich**, H Reich**, P Ling^#^, M Romano^#^

*University of Miami, Miami, FL:* A Fornoni^*^, C Bidot^#^

*University of Michigan, Ann Arbor, MI:* M Kretzler^*^, D Gipson*, A Williams^#^, C Klida^#^

*University of North Carolina, Chapel Hill, NC:* V Derebail^*^, K Gibson^*^, E Cole^#^, J Ormond-Foster^#^

*University of Pennsylvania, Philadelphia, PA:* L Holzman^*^, K Meyers**, K Kallem^#^, A Swenson^#^

*University of Texas Southwestern, Dallas, TX:* K Sambandam^*^, Z Wang^#^, M Rogers^#^

*University of Washington, Seattle, WA:* A Jefferson^*^, S Hingorani**, K Tuttle**^§^, M Bray ^#^, E Pao^#^, A Cooper^#§^

*Wake Forest University Baptist Health, Winston-Salem, NC:* JJ Lin*, Stefanie Baker^#^

*Data Analysis and Coordinating Center*: M Kretzler*, L Barisoni**, J Bixler, H Desmond, S Eddy, D Fermin, C Gadegbeku**, B Gillespie**, D Gipson**, L Holzman**, V Kurtz, M Larkina, S Li, S Li, CC Lienczewski, J Liu, T Mainieri, L Mariani**, M Sampson**, J Sedor**, A Smith, A Williams, J Zee.

*Digital Pathology Committee*: Carmen Avila-Casado (University Health Network, Toronto), Serena Bagnasco (Johns Hopkins University), Joseph Gaut (Washington University in St Louis), Stephen Hewitt (National Cancer Institute), Jeff Hodgin (University of Michigan), Kevin Lemley (Children’s Hospital of Los Angeles), Laura Mariani (University of Michigan), Matthew Palmer (University of Pennsylvania), Avi Rosenberg (Johns Hopkins University), Virginie Royal (University of Montreal), David Thomas (University of Miami), Jarcy Zee (University of Pennsylvania) Co-Chairs: Laura Barisoni (Duke University) and Cynthia Nast (Cedar Sinai).

*Principal Investigator; **Co-investigator^; #^Study Coordinator

^§^Providence Medical Research Center, Spokane, WA

## Funding Acknowledgements

MCM wishes to acknowledge funding from RO1-DK122164, American Heart Association (AHASDG25870018), and philanthropy from Nina and Homer Eaton.

The Nephrotic Syndrome Study Network Consortium (NEPTUNE), U54□DK□083912, is a part of the National Institutes of Health (NIH) Rare Disease Clinical Research Network (RDCRN), supported through a collaboration between the Office of Rare Diseases Research, National Center for Advancing Translational Sciences and the National Institute of Diabetes, Digestive, and Kidney Diseases. Additional funding and/or programmatic support for this project has also been provided by the University of Michigan, the NephCure Kidney International and the Halpin Foundation.

## References

1. Menon MC, Chuang PY, and He CJ. The glomerular filtration barrier: components and crosstalk. Int J Nephrol. 2012;2012:749010.

2. Maas RJ, Deegens JK, Smeets B, Moeller MJ, and Wetzels JF. Minimal change disease and idiopathic FSGS: manifestations of the same disease. Nat Rev Nephrol. 2016;12(12):768–76.

3. Fufaa GD, Weil EJ, Lemley KV, Knowler WC, Brosius FC, 3rd, Yee B, et al. Structural Predictors of Loss of Renal Function in American Indians with Type 2 Diabetes. Clin J Am Soc Nephrol. 2016;11(2):254–61.

4. Puelles VG, van der Wolde JW, Wanner N, Scheppach MW, Cullen-McEwen LA, Bork T, et al. mTOR-mediated podocyte hypertrophy regulates glomerular integrity in mice and humans. JCI Insight. 2019;4(18).

5. Lemley KV, Bagnasco SM, Nast CC, Barisoni L, Conway CM, Hewitt SM, et al. Morphometry Predicts Early GFR Change in Primary Proteinuric Glomerulopathies: A Longitudinal Cohort Study Using Generalized Estimating Equations. PLoS One. 2016;11(6):e0157148.

6. Romoli S, Angelotti ML, Antonelli G, Kumar Vr S, Mulay SR, Desai J, et al. CXCL12 blockade preferentially regenerates lost podocytes in cortical nephrons by targeting an intrinsic podocyte-progenitor feedback mechanism. Kidney Int. 2018;94(6):1111–26.

7. Velez JCQ, Arif E, Rodgers J, Hicks MP, Arthur JM, Nihalani D, et al. Deficiency of the Angiotensinase Aminopeptidase A Increases Susceptibility to Glomerular Injury. J Am Soc Nephrol. 2017;28(7):2119–32.

8. Nishizono R, Kikuchi M, Wang SQ, Chowdhury M, Nair V, Hartman J, et al. FSGS as an Adaptive Response to Growth-Induced Podocyte Stress. J Am Soc Nephrol. 2017;28(10):2931–45.

9. Menon MC, Chuang PY, Li Z, Wei C, Zhang W, Luan Y, et al. Intronic locus determines SHROOM3 expression and potentiates renal allograft fibrosis. J Clin Invest. 2014.

10. Wei C, Banu K, Garzon F, Basgen JM, Philippe N, Yi Z, et al. SHROOM3-FYN Interaction Regulates Nephrin Phosphorylation and Affects Albuminuria in Allografts. J Am Soc Nephrol. 2018;29(11):2641–57.

11. Prokop JW, Yeo NC, Ottmann C, Chhetri SB, Florus KL, Ross EJ, et al. Characterization of Coding/Noncoding Variants for SHROOM3 in Patients with CKD. J Am Soc Nephrol. 2018.

12. Yeo NC, O’Meara CC, Bonomo JA, Veth KN, Tomar R, Flister MJ, et al. Shroom3 contributes to the maintenance of the glomerular filtration barrier integrity. Genome Res. 2014.

13. Verma R, Wharram B, Kovari I, Kunkel R, Nihalani D, Wary KK, et al. Fyn binds to and phosphorylates the kidney slit diaphragm component Nephrin. J Biol Chem. 2003;278(23):20716–23.

14. Audard V, Zhang SY, Copie-Bergman C, Rucker-Martin C, Ory V, Candelier M, et al. Occurrence of minimal change nephrotic syndrome in classical Hodgkin lymphoma is closely related to the induction of c-mip in Hodgkin-Reed Sternberg cells and podocytes. Blood. 2010;115(18):3756–62.

15. Zhang SY, Kamal M, Dahan K, Pawlak A, Ory V, Desvaux D, et al. c-mip impairs podocyte proximal signaling and induces heavy proteinuria. Sci Signal. 2010;3(122):ra39.

16. Gadegbeku CA, Gipson DS, Holzman LB, Ojo AO, Song PX, Barisoni L, et al. Design of the Nephrotic Syndrome Study Network (NEPTUNE) to evaluate primary glomerular nephropathy by a multidisciplinary approach. Kidney Int. 2013;83(4):749–56.

17. Menon MC, Chuang PY, Li Z, Wei C, Zhang W, Luan Y, et al. Intronic locus determines SHROOM3 expression and potentiates renal allograft fibrosis. J Clin Invest. 2015;125(1):208–21.

18. Premsrirut PK, Dow LE, Kim SY, Camiolo M, Malone CD, Miething C, et al. A rapid and scalable system for studying gene function in mice using conditional RNA interference. Cell. 2011;145(1):145–58.

19. Basgen JM, and Sobin C. Early chronic low-level lead exposure produces glomerular hypertrophy in young C57BL/6J mice. Toxicol Lett. 2014;225(1):48–56.

20. Lin X, Suh JH, Go G, and Miner JH. Feasibility of repairing glomerular basement membrane defects in Alport syndrome. J Am Soc Nephrol. 2014;25(4):687–92.

21. Topham PS, Haydar SA, Kuphal R, Lightfoot JD, and Salant DJ. Complement-mediated injury reversibly disrupts glomerular epithelial cell actin microfilaments and focal adhesions. Kidney Int. 1999;55(5):1763–75.

22. Yamada E, Pessin JE, Kurland IJ, Schwartz GJ, and Bastie CC. Fyn-dependent regulation of energy expenditure and body weight is mediated by tyrosine phosphorylation of LKB1. Cell Metab. 2010;11(2):113–24.

23. Johanns M, Pyr Dit Ruys S, Houddane A, Vertommen D, Herinckx G, Hue L, et al. Direct and indirect activation of eukaryotic elongation factor 2 kinase by AMP-activated protein kinase. Cell Signal. 2017;36:212–21.

24. Cao Y, Bojjireddy N, Kim M, Li T, Zhai P, Nagarajan N, et al. Activation of gamma2-AMPK Suppresses Ribosome Biogenesis and Protects Against Myocardial Ischemia/Reperfusion Injury. Circ Res. 2017;121(10):1182–91.

25. Riediger F, Quack I, Qadri F, Hartleben B, Park JK, Potthoff SA, et al. Prorenin receptor is essential for podocyte autophagy and survival. J Am Soc Nephrol. 2011;22(12):2193–202.

26. Narbonne P, and Roy R. Caenorhabditis elegans dauers need LKB1/AMPK to ration lipid reserves and ensure long-term survival. Nature. 2009;457(7226):210–4.

27. Steinberg GR, and Kemp BE. AMPK in Health and Disease. Physiol Rev. 2009;89(3):1025–78.

28. Longo VD, Antebi A, Bartke A, Barzilai N, Brown-Borg HM, Caruso C, et al. Interventions to Slow Aging in Humans: Are We Ready? Aging Cell. 2015;14(4):497–510.

29. Takiar V, Nishio S, Seo-Mayer P, King JD, Jr., Li H, Zhang L, et al. Activating AMP-activated protein kinase (AMPK) slows renal cystogenesis. Proc Natl Acad Sci U S A. 2011;108(6):2462–7.

30. Abdulrahman RM, Boon MR, Sips HC, Guigas B, Rensen PC, Smit JW, et al. Impact of Metformin and compound C on NIS expression and iodine uptake in vitro and in vivo: a role for CRE in AMPK modulation of thyroid function. Thyroid. 2014;24(1):78–87.

31. Zhou G, Myers R, Li Y, Chen Y, Shen X, Fenyk-Melody J, et al. Role of AMP-activated protein kinase in mechanism of metformin action. J Clin Invest. 2001;108(8):1167–74.

32. McCullough LD, Zeng Z, Li H, Landree LE, McFadden J, and Ronnett GV. Pharmacological inhibition of AMP-activated protein kinase provides neuroprotection in stroke. J Biol Chem. 2005;280(21):20493–502.

33. Wu CI, Lu YY, Chen YC, Lin FZ, Huang JH, Lin YK, et al. The AMP-activated protein kinase modulates hypothermia-induced J wave. Eur J Clin Invest. 2020;50(6):e13247.

34. Wharram BL, Goyal M, Wiggins JE, Sanden SK, Hussain S, Filipiak WE, et al. Podocyte depletion causes glomerulosclerosis: diphtheria toxin-induced podocyte depletion in rats expressing human diphtheria toxin receptor transgene. J Am Soc Nephrol. 2005;16(10):2941–52.

35. Yang Y, Hodgin JB, Afshinnia F, Wang SQ, Wickman L, Chowdhury M, et al. The two kidney to one kidney transition and transplant glomerulopathy: a podocyte perspective. J Am Soc Nephrol. 2015;26(6):1450–65.

36. Kikuchi M, Wickman L, Hodgin JB, and Wiggins RC. Podometrics as a Potential Clinical Tool for Glomerular Disease Management. Semin Nephrol. 2015;35(3):245–55.

37. Carling D. AMPK signalling in health and disease. Curr Opin Cell Biol. 2017;45:31–7.

38. Bork T, Liang W, Yamahara K, Lee P, Tian Z, Liu S, et al. Podocytes maintain high basal levels of autophagy independent of mtor signaling. Autophagy. 2019:1–17.

39. Egan DF, Shackelford DB, Mihaylova MM, Gelino S, Kohnz RA, Mair W, et al. Phosphorylation of ULK1 (hATG1) by AMP-activated protein kinase connects energy sensing to mitophagy. Science. 2011;331(6016):456–61.

40. Yi M, Zhang L, Liu Y, Livingston MJ, Chen JK, Nahman NS, Jr., et al. Autophagy is activated to protect against podocyte injury in adriamycin-induced nephropathy. Am J Physiol Renal Physiol. 2017;313(1):F74–F84.

41. Decleves AE, Mathew AV, Cunard R, and Sharma K. AMPK mediates the initiation of kidney disease induced by a high-fat diet. J Am Soc Nephrol. 2011;22(10):1846–55.

42. Dugan LL, You YH, Ali SS, Diamond-Stanic M, Miyamoto S, DeCleves AE, et al. AMPK dysregulation promotes diabetes-related reduction of superoxide and mitochondrial function. J Clin Invest. 2013;123(11):4888–99.

43. Satriano J, Sharma K, Blantz RC, and Deng A. Induction of AMPK activity corrects early pathophysiological alterations in the subtotal nephrectomy model of chronic kidney disease. Am J Physiol Renal Physiol. 2013;305(5):F727–33.

44. Borges CM, Fujihara CK, Malheiros D, de Avila VF, Formigari GP, and Lopes de Faria JB. Metformin arrests the progression of established kidney disease in the subtotal nephrectomy model of chronic kidney disease. Am J Physiol Renal Physiol. 2020;318(5):F1229–F36.

45. Steinberg GR, and Carling D. AMP-activated protein kinase: the current landscape for drug development. Nat Rev Drug Discov. 2019;18(7):527–51.

46. Bhaskar Das DPW. LKB1-AMPK ACTIVATORS FOR THERAPEUTIC USE IN POLYCYSTIC KIDNEY DISEASE. https://pubchem.ncbi.nlm.nih.gov/patent/US2017334892#section=Patent-Submission-Date.

47. Yamada E, Okada S, Bastie CC, Vatish M, Nakajima Y, Shibusawa R, et al. Fyn phosphorylates AMPK to inhibit AMPK activity and AMP-dependent activation of autophagy. Oncotarget. 2016;7(46):74612–29.

48. Kir S, Komaba H, Garcia AP, Economopoulos KP, Liu W, Lanske B, et al. PTH/PTHrP Receptor Mediates Cachexia in Models of Kidney Failure and Cancer. Cell Metab. 2016;23(2):315–23.

49. Bachmanov AA, Reed DR, Beauchamp GK, and Tordoff MG. Food intake, water intake, and drinking spout side preference of 28 mouse strains. Behav Genet. 2002;32(6):435–43.

50. Er W. Stereological Methods. London: Academic Press; 1979:44–5.

51. Bai XY, and Basgen JM. Podocyte number in the maturing rat kidney. Am J Nephrol. 2011;33(1):91–6.

