## Supplemental Figures for "AMP-Kinase mediates regulation of glomerular volume and podocyte survival"

**Figure S1**

A B

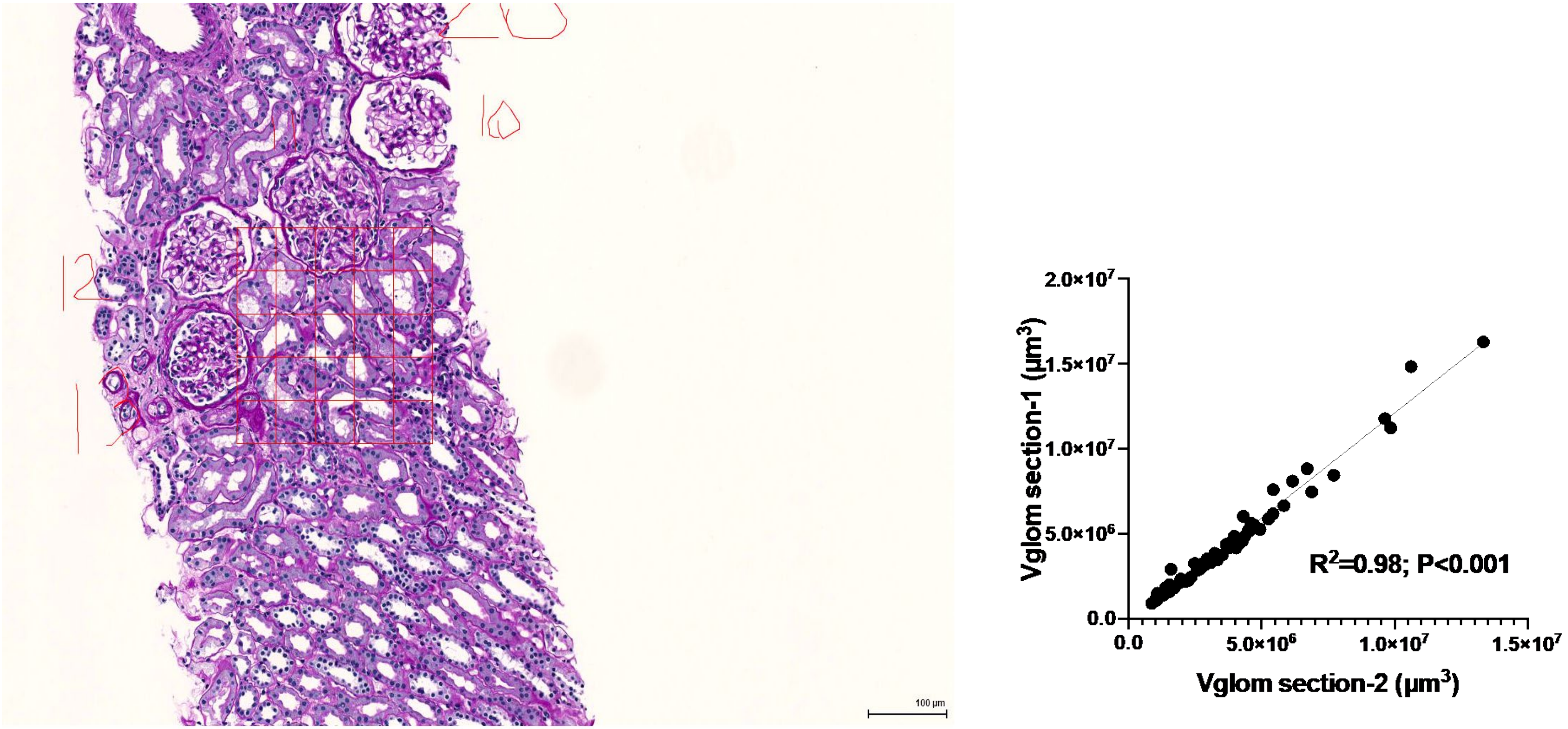


**Figure S1. Glomerular morphometry shows significantly lower Vglom in MCD vs FSGS cases. (1A)** Aperio-scanned image of representative NS biopsy from NEPTUNE study. Glomerular volumes were calculated from the Weibel-Gomez method from area cross sections of glomeruli measured by planimetry on PAS images. Weibel-Gomez method: Vglom = 1.38 x Area$^{3/2}$ where Vglom is the glomerular volume, 1.38 is the assumed shape factor for the glomeruli, and Area is the average cross-sectional tuft area of the glomerular profiles measured. Image adapted from Lemley et al, PLOSone, 2016 (5). **(1B)** Correlation plot of Vglom (in μm$^3$) from two random sections obtained with the same biopsy ($R^2$=0.98; P<0.001; Spearman R). [PAS= periodic acid Schiff]

**Figure S2A-B**

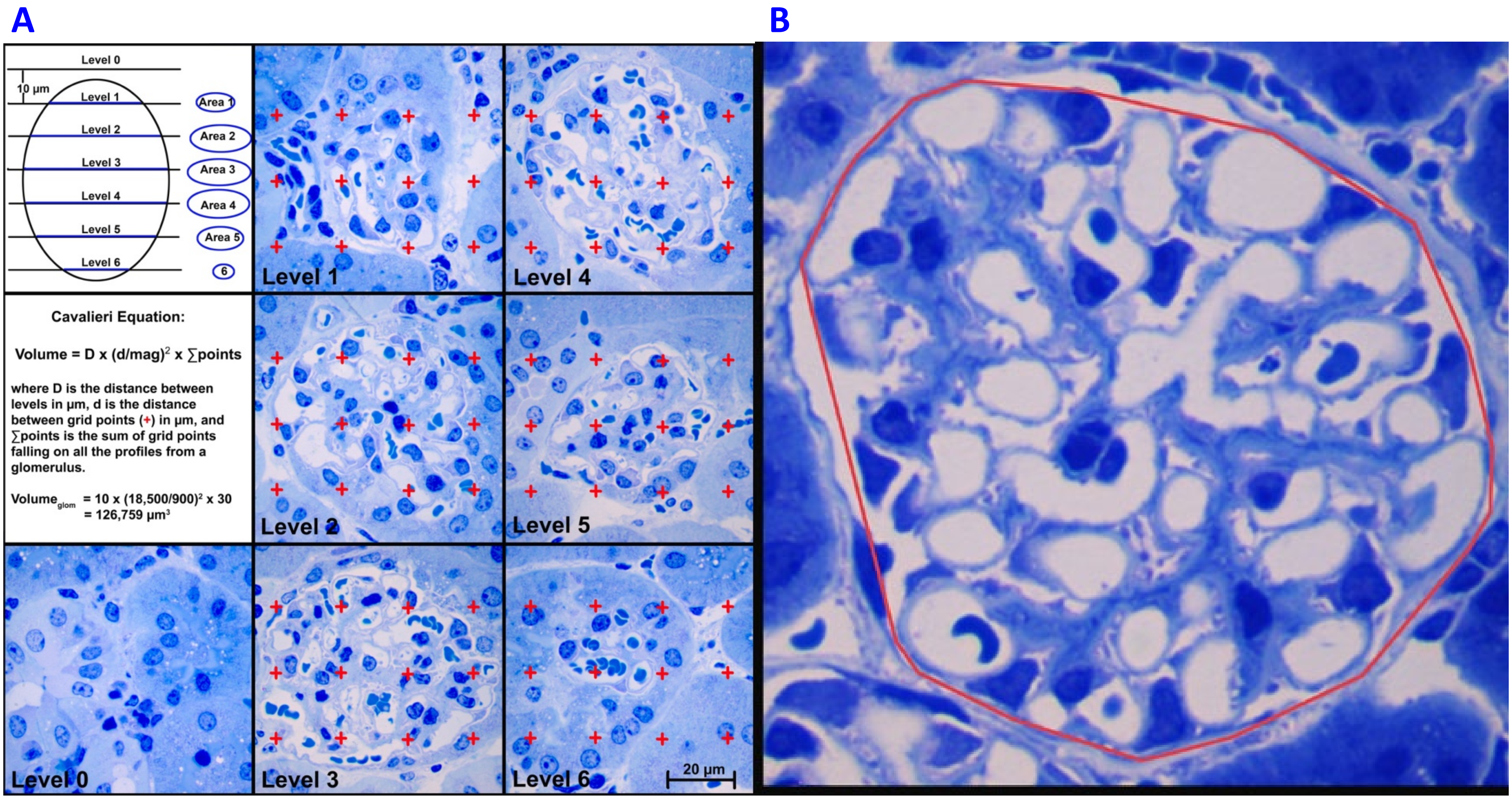


**Figure S2A-B. Global or Podocyte specific Shroom3 knockdown reduced glomerular and podocyte volume: (2A)** Demonstration of Cavalieri method to measure Glomerular volume. Upper left panel demonstrates the sampling scheme for the Cavalieri method to measure volume of an arbitrary particle. Middle left panel shows the Cavalieri equation. Remaining panels are images through a sample glomerulus with grid points superimposed over the toluidine blue epon sections. **(2B**) Representative image used for Vglom component analyses. Glomerular profile is defined by a minimal polygon around the glomerular profile in image **(see 2C & 2D & methods)**

**Figure S2C-D**

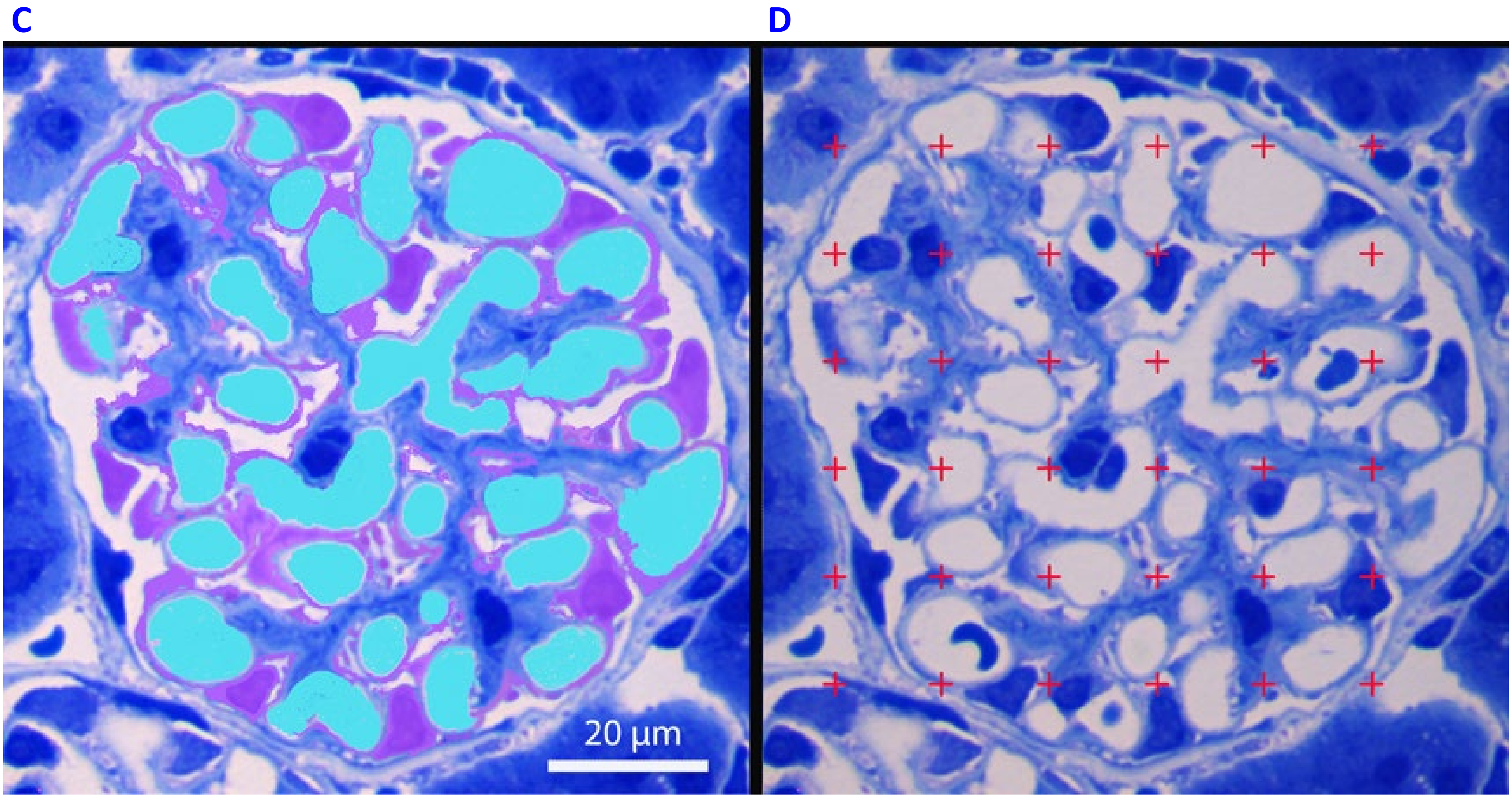


**Figure S2C-D. Global or Podocyte specific Shroom3 knockdown reduced glomerular and podocyte volume (continued):** Representative image used for Vglom component analyses are shown. Glomerular profile is defined by a minimal polygon around the glomerular profile in image as in **2B**. **(2C)** Each glomerular profile is divided into four Vglom components as demarcated in pseudocolor – podocytes (purple), Capillary space+endothelial cell (cyan), mesangium (original blue), and other (white within polygon). **(2D)** A counting grid is randomly placed over the glomerular profile and number of points falling on each component is counted. 10 gloms &~ 600 points were counted per kidney.

Formula: ***Volume density of Component X=Σ points on X/ Σ points on Glomerulus***

Figure S2E-J

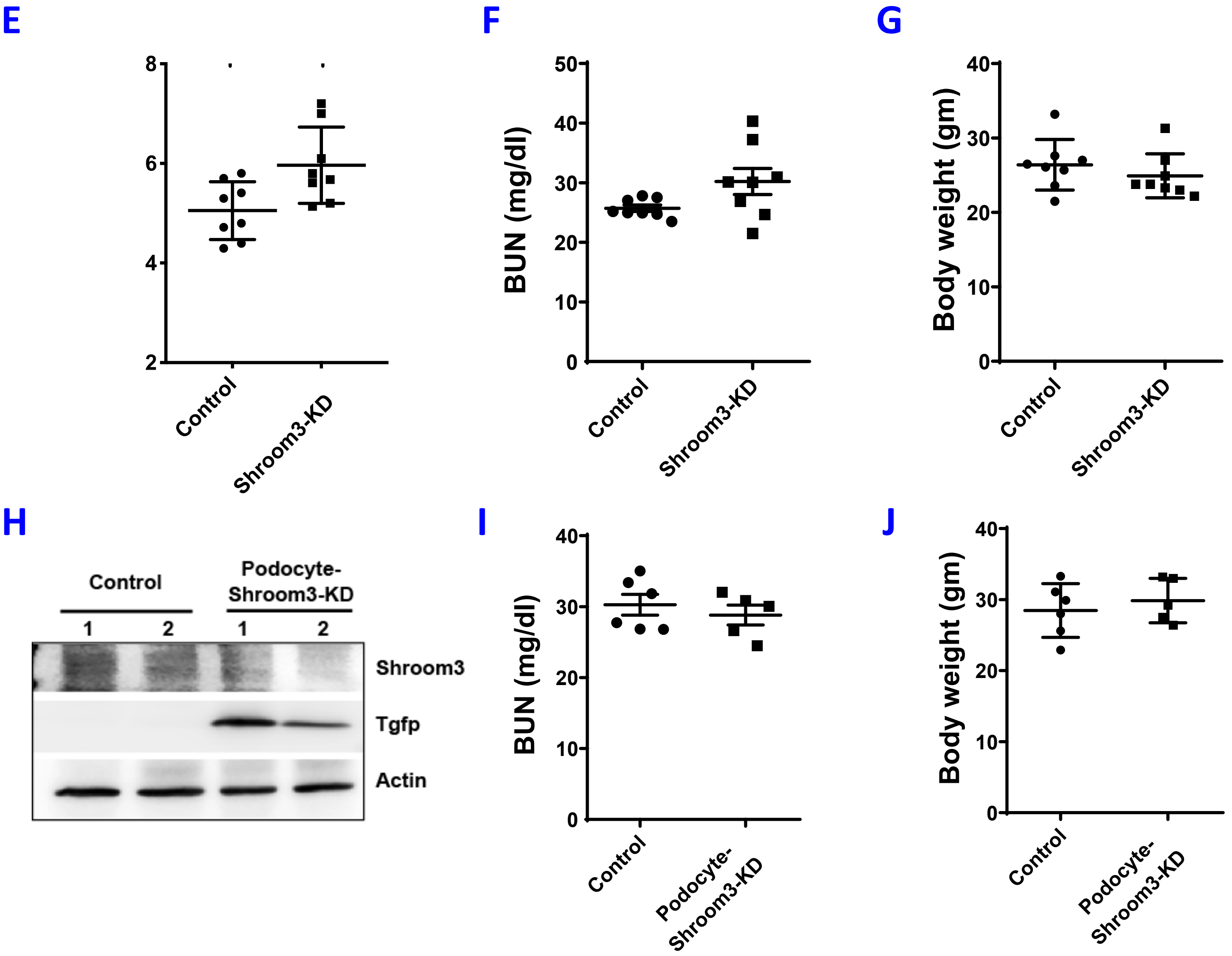


**Figure S2E-J. Global or Podocyte specific Shroom3 knockdown reduced glomerular and podocyte volume:** Control & Shroom3 KD mice were DOX fed for 6 weeks (n=8 each). Dot plots compare **(2E)** mean podocyte nuclear density (per $\mu m^3$ Vglom) , **(2F)** mean blood urea nitrogen levels (mg/dl), and **(2G)** body weights (gms) . Similarly, Control and Podocyte-Shroom3-KD mice were DOX fed for 6-weeks (n=6 vs 5). **(2H)** Representative WBs of Glomerular lysates (n=2 each group) demonstrate immunoblotting for Shroom3, TurboGFP, and Actin. Dot-plots compare **(2I)** mean blood urea nitrogen levels (mg/dl), and **(2J)** body weights (gms) of control and Podocyte Shroom3-KD mice [Line/Whiskers=Mean/SEM; *= P<0.05].

**Figure S3**

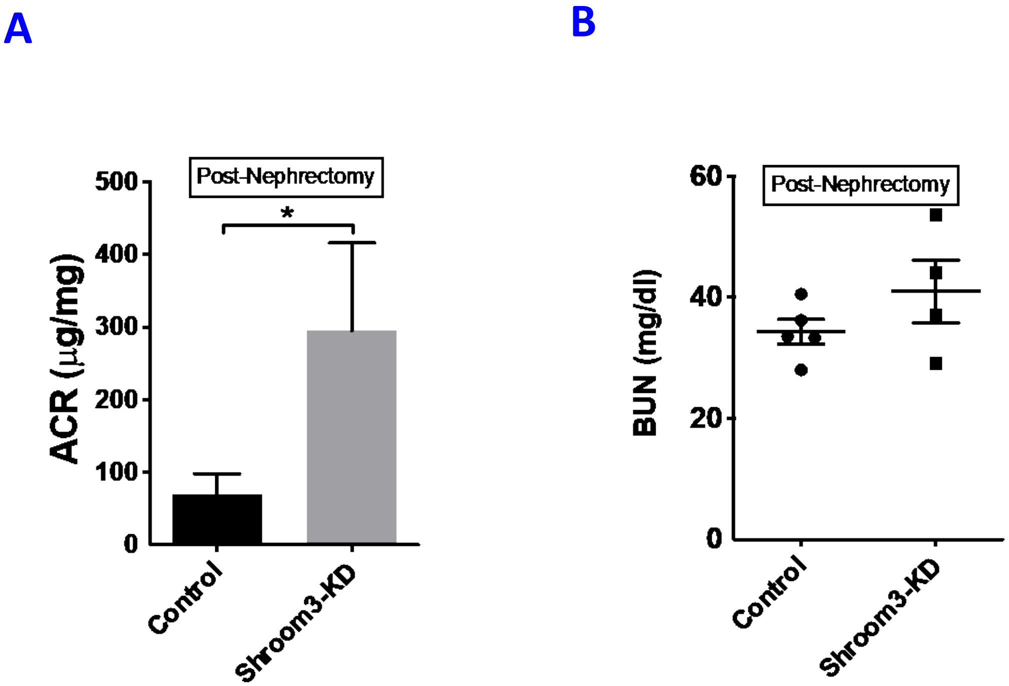


**Figure S3. Shroom3 knockdown restricted glomerular hypertrophy post-unilateral nephrectomy:** Control & Shroom3-KD mice underwent unilateral nephrectomy (n=5 vs 4). Remnant kidney was evaluated at 1-week post nephrectomy. Bar graph show Albumin: creatinine ratio (µg/mg) **(3A),** and dot plot show BUN (mg/dl) **(3B)** at 1 week post nephrectomy**.** Dot [Line/Whiskers=Mean/SEM; *= P<0.05].

**Figure S4**

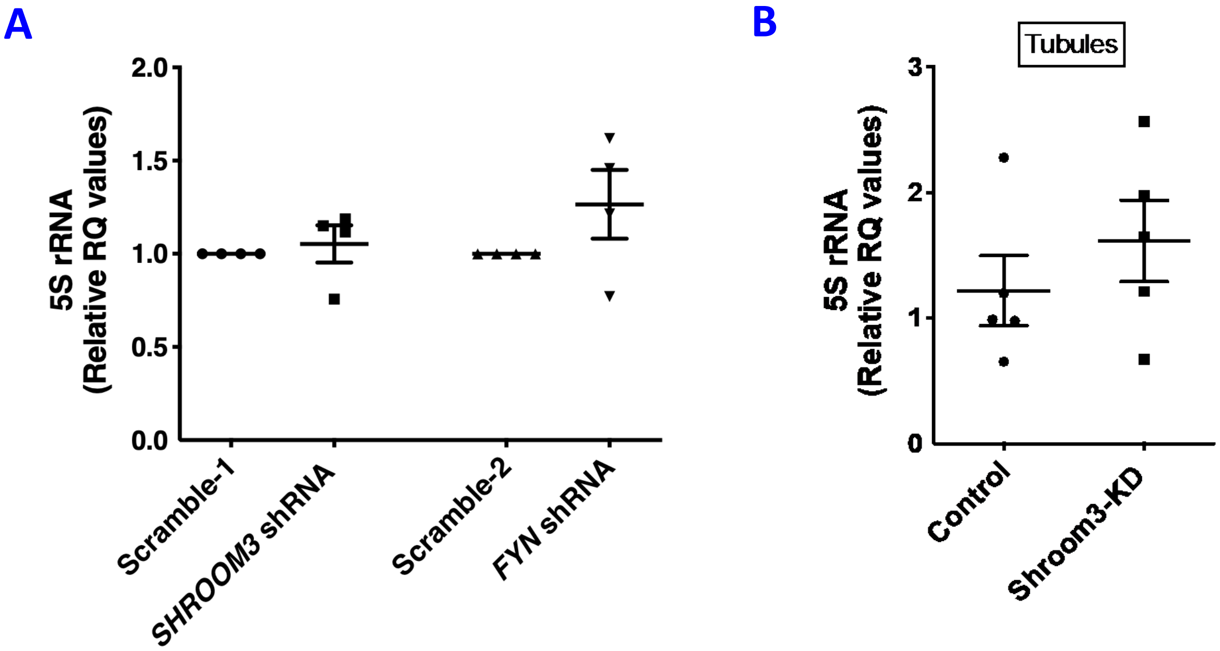


**Figure S4. Shroom3 knockdown reduces cellular protein content and RNA biogenesis *in vitro* and *in vivo* mediated via FYN: (4A)** Puromycin-selectable, stable Shroom3 and Fyn knockdown podocytes were generated using lentiviral shRNA infection (Scramble-1 and Scramble-2 are respective scramble-sequence infected controls). Stable podocytes were differentiated (>7 days) in collagen coated plates. In SHROOM3- & FYN-shRNA podocyte lines, dot plots show paired comparisons (n=4 sets) of copy numbers of 5S subunit of ribosomal RNA (5S rRNA) normalized to Actin, vs Scramble infected podocyte lines **(4B)** Dot plots show compare copy numbers of 5S rRNA normalized to Actin, in tubular RNA extracts (n=5 vs 5 Control vs Shroom3-KD mice). [Line/Whiskers=Mean/SEM; compared by paired t-test (A) and unpaired t-test (B); RQ=Relative quantity].

**Figure S5A-D**

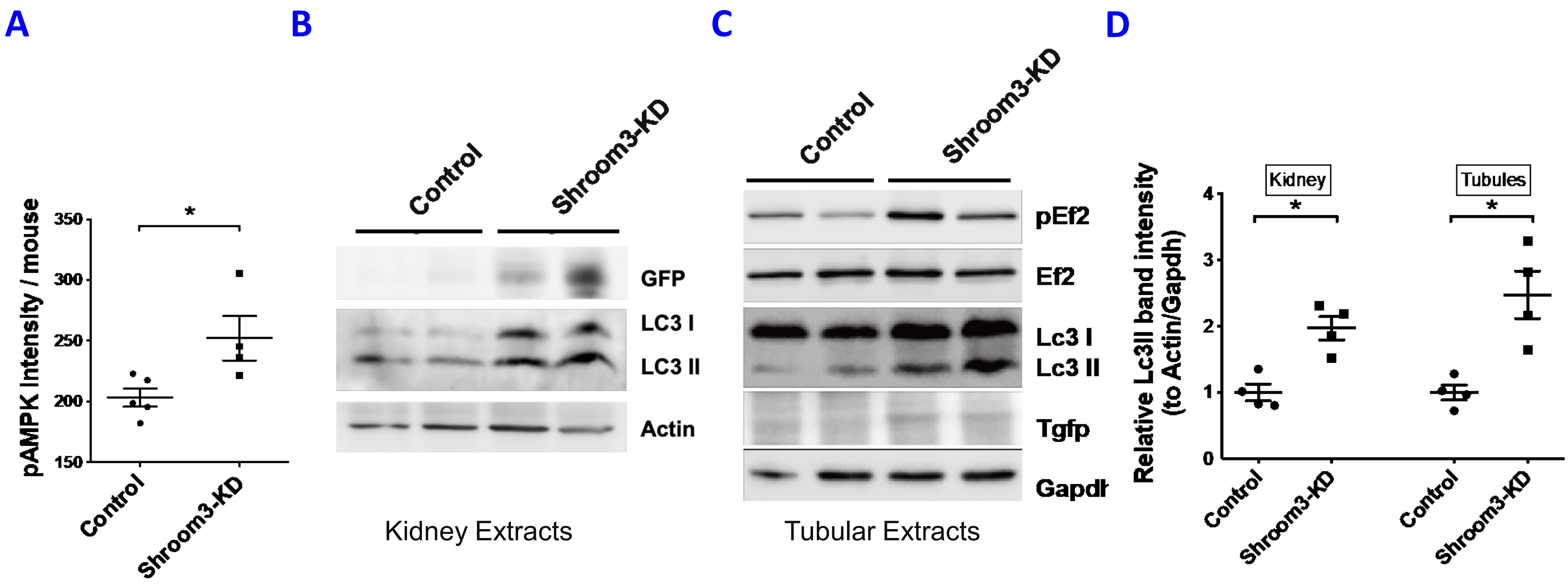


**Figure S5A-D. Shroom3- or Fyn knockdown increase cellular AMPK activation: (5A)** Dot plots show quantification of intensity of Phosphorylated AMPK/per glomerular outline per group (depicted per animal). Representative images of immunoblots from Control vs Shroom3-KD (n=2 each) showing **(5B)** GFP, Lc3 I & II and Actin bands in kidney extracts and **(5C)** showing immunoblots from kidney tubular extracts from Control vs Shroom3-KD (n=2 each) showing pEf2, Ef2, Lc3 I & II, Tgfp and Gapdh bands. **(5D)** Dot plots show the respective relative band intensity of Lc3-II in Control vs Shroom3-KD lysates (normalized to Actin/Gapdh; n=4 mice). [Line/Whiskers = Mean/ SEM; *= P<0.05; unpaired t-test ].

### Figure S5E-F

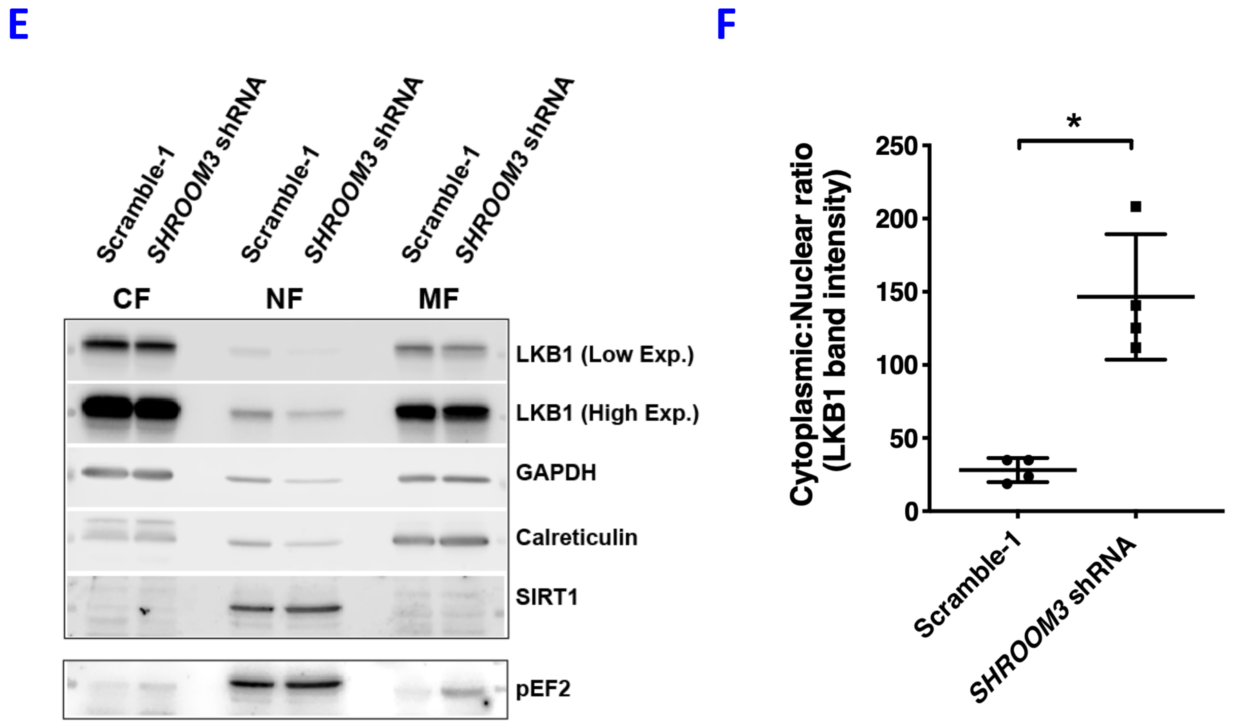


**Figure-S5E-F. Shroom3- or Fyn knockdown increase cellular AMPK activation:** Reduced nuclear retention of LKB1 with FYN inactivation or knockdown results in LKB1-mediated AMPK activation in podocytes. **(5E)** Representative WBs of lysates obtained after sub-cellular protein fractionation (CF/NF/MF=Cytoplasmic-/Nuclear-/Membrane Fractions, respectively) from Scramble-1, *SHROOM3*-shRNA podocytes probed for LKB1, Phosphorylated EF2, as well as GAPDH, Calreticulin, SIRT1 used as fractionation controls for CF, MF and NF, respectively. **(5F)** Dot plots show the respective relative band intensity of LKB1 (CF:NF ratio; n=4 sets). [Line/Whiskers=Mean/SEM; unpaired t-test; *= P<0.05].

**Figure S5G**

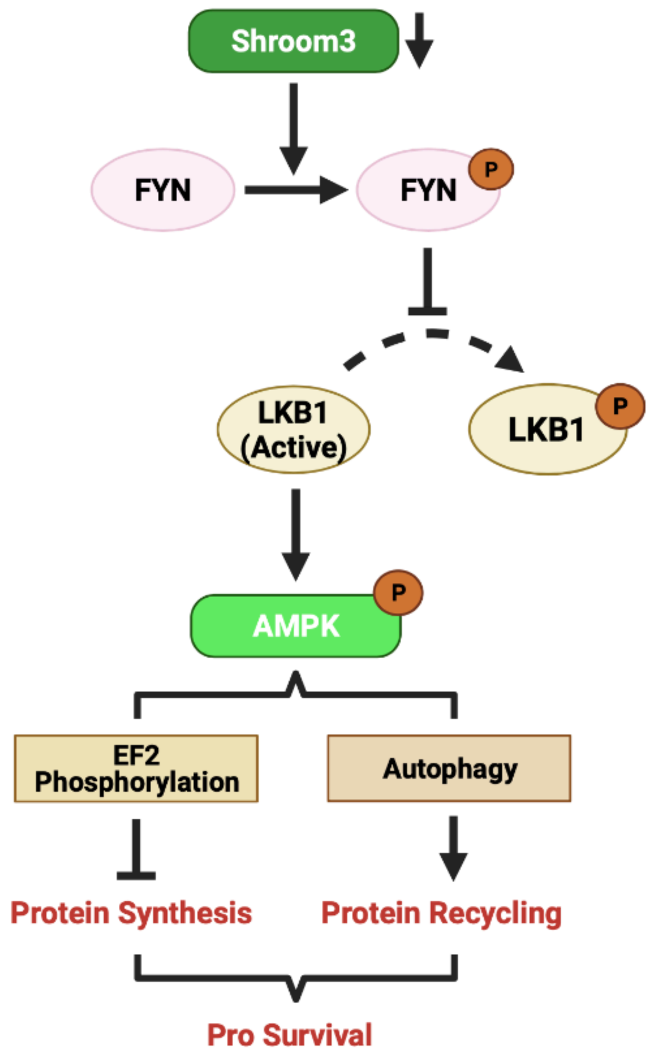


**Figure-S5G. Shroom3- or Fyn knockdown increase cellular AMPK activation:** Signaling schema summarizes AMPK activation in podocytes with Shroom3 knockdown mediated via LKB1 cytoplasmic redistribution downstream of Shroom3-Fyn axis. AMPK activation regulates cell and glomerular size by engaging downstream mechanisms - reduced anabolism and increased autophagy.

**Figure S6**

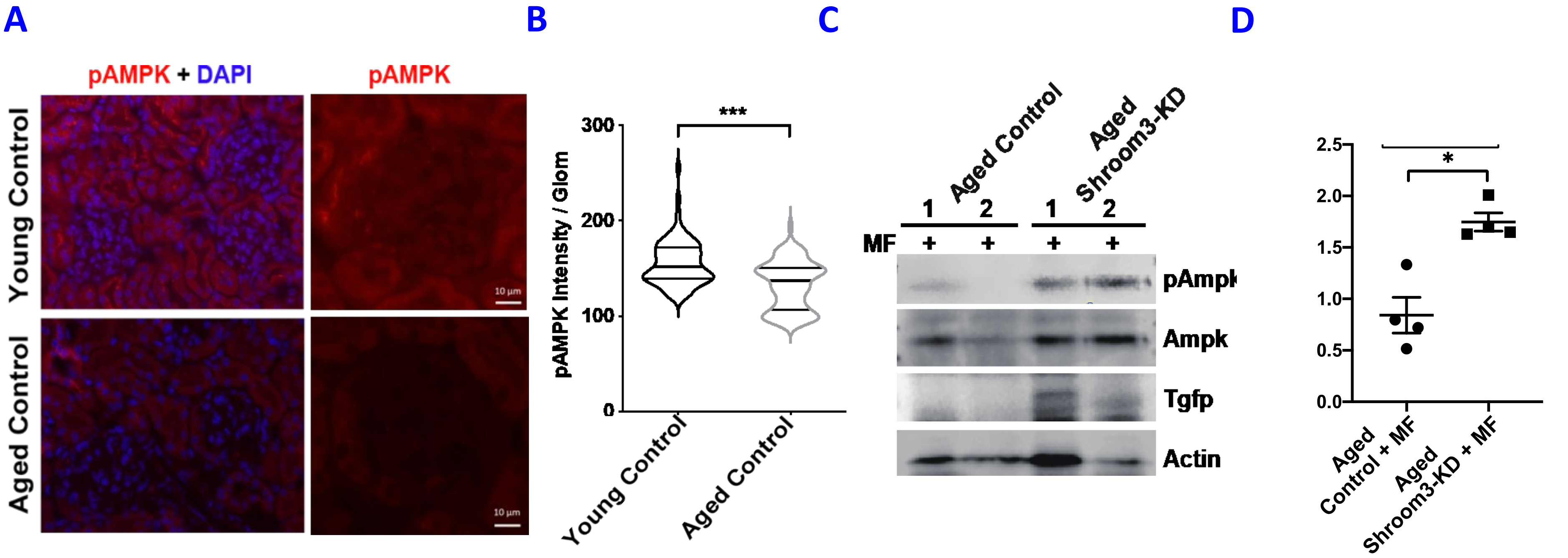


**Figure S6. AMPK activation reduces Vglom and mitigates podocytopenia in aged Shroom3 knockdown mice with podocyte FPE:** As described previously, Control & Shroom3-KD mice were aged > 1-year and DOX feeding for 6 weeks. In subsequent experiments, at week-2, Metformin-water (MF) was added. **(6A)** Representative immunofluorescence images showing Phosphorylated AMPK/WT1/DAPI in young Controls, & Aged Shroom3-KD mice **(6B)** Violin -plots show quantification of intensity of Phosphorylated AMPK/per glomerular outline per group (depicted per glomerulus; 30 gloms/animal; n=5 each). (**6C)** Representative images of immunoblots from Aged Controls vs Shroom3-KD + MF (n=2 each) showing Phospho-Ampk, Ampk, Tgfp and Actin **(6D)** Dot plots show the respective relative band intensity of pAmpk:Ampk (to Actin; n=4) [Line/Whiskers=Mean/SEM; unpaired t-test; *= P<0.05, ***= P<0.001].

**Figure S7**

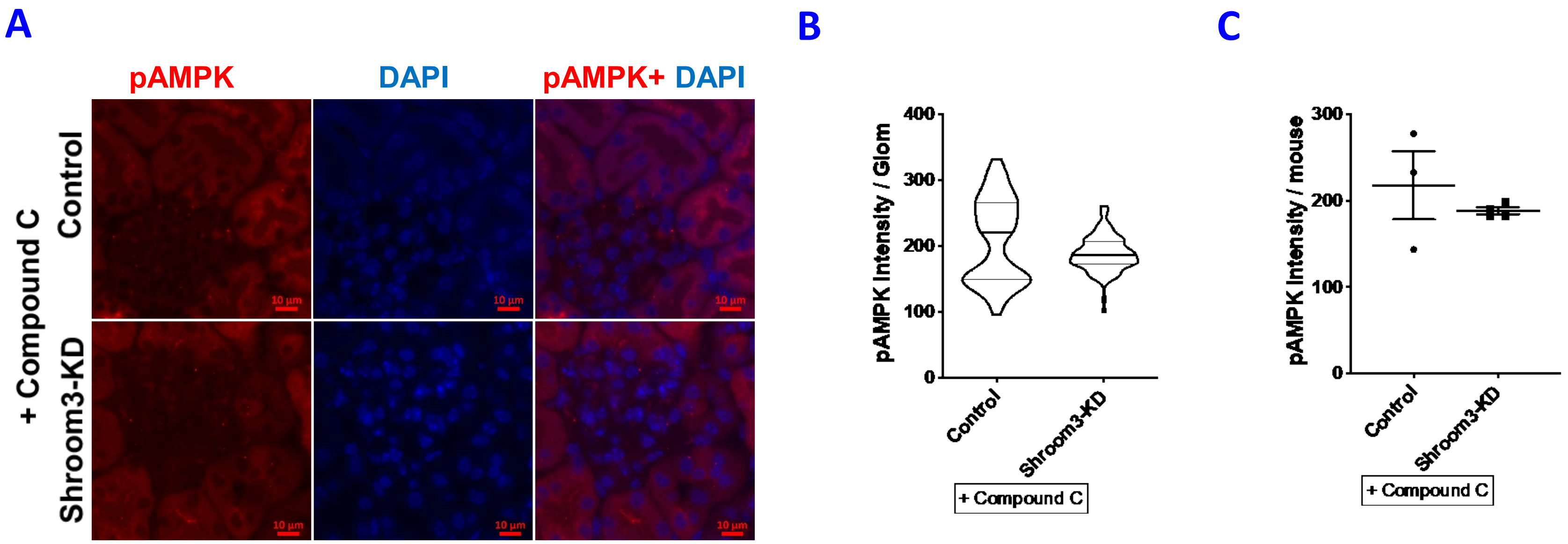


**Figure S7. AMPK inhibition reverses Vglom reduction and promotes podocytopenia in Shroom3 knockdown mice:** Control vs Shroom3-KD mice (~8 weeks) were DOX fed for 8 weeks and administered Compound C at week 5 **(7A)** Representative immunofluorescence images (40X) show Phosphorylated AMPK/DAPI in Control vs (upper row) & Shroom3-KD mice (lower row). **(7B)** Violin -plots show quantification of intensity of Phosphorylated AMPK/per glomerular outline per group (depicted per glomerulus; 30 gloms/animal). **(7C)** Dot plots quantify intensity of Phosphorylated AMPK/per glomerular outline per animal (n=3 vs 4), [Line/Whiskers=Mean/SEM; unpaired t-test]

Figure S8A-D

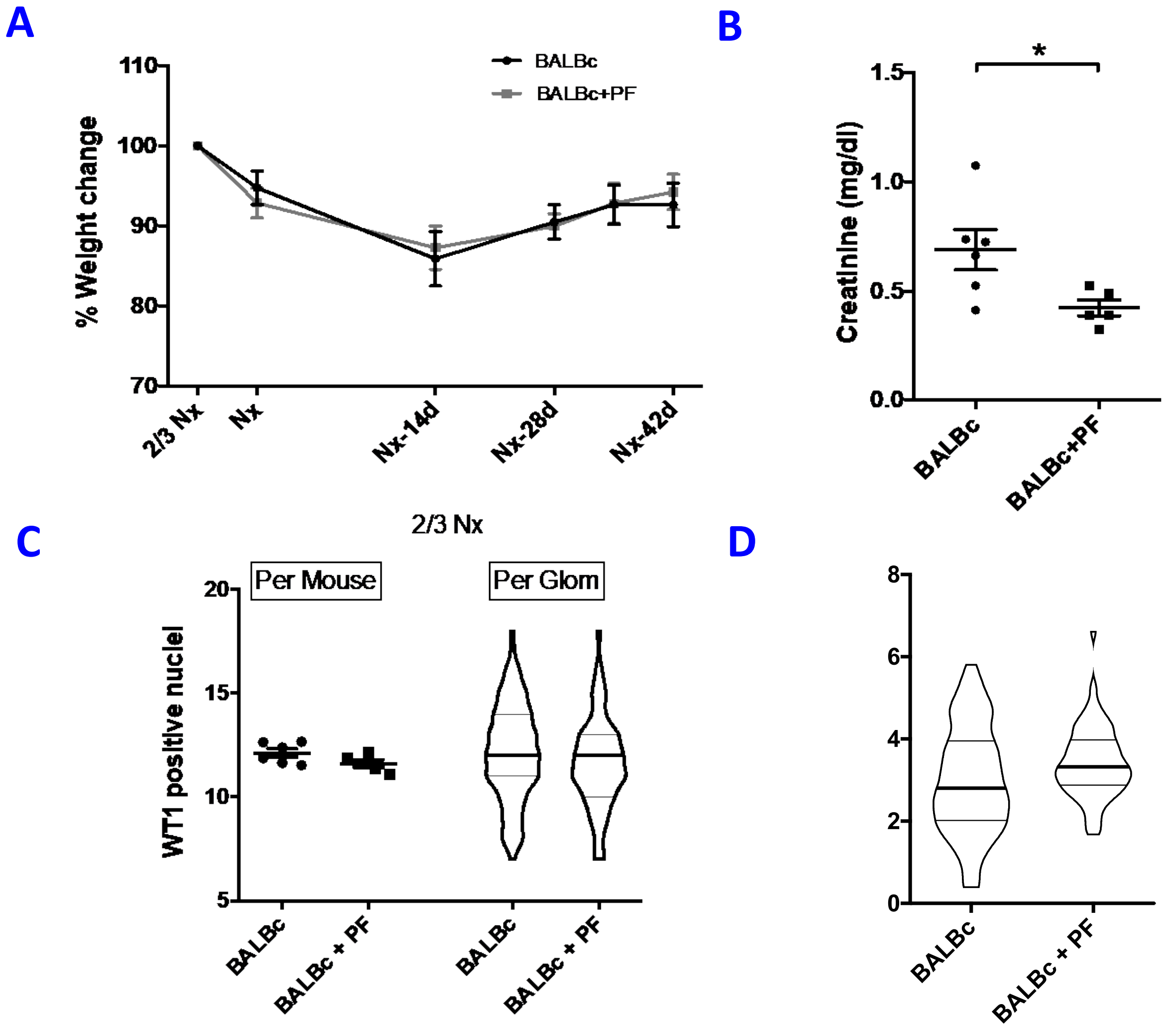


**Figure S8A-D. AMPK-activation reduces glomerular volume and preserves podocyte numbers in nephron loss-induced glomerular hypertrophy:** Adult BALBc mice (near 8-wks) underwent 2/3$^{rd}$ nephrectomy (2/3 Nx), followed by contralateral nephrectomy (Nx) 1 week later, and were followed for 6 more weeks when the remaining kidney tissue (1/6$^{th}$ remnant) was harvested. Experimental animals were gavaged with AMPK-activator PF06409577 (3 doses/week; BALBc (n=6) vs BALBc+PF (n=5)). **(8A)** Lines show mean weight trends (as percentage of initial weight). **(8B)** Dot-plots compare Creatinine (mg/dl) in these groups at 6 weeks**. (8C)** Dot plots compare mean podocyte numbers/glomerulus/animal while Violin-plots (Line at median) depict distribution of mean podocytes/ glomerulus/ group (30 glomerular profiles/mouse) in 2/3 Nx kidneys obtained following first surgery. **(8D)** Violin-plots show distribution of podocyte nuclear density per glomerulus (per μm$^3$ Vglom) in each group (Line at median; n=10 glomeruli/mouse**).** [Line/Whiskers = Mean/SEM; unpaired t-test; *= P<0.05; WT1= Wilm's Tumor-1 protein].

**Figure S8 E-I**

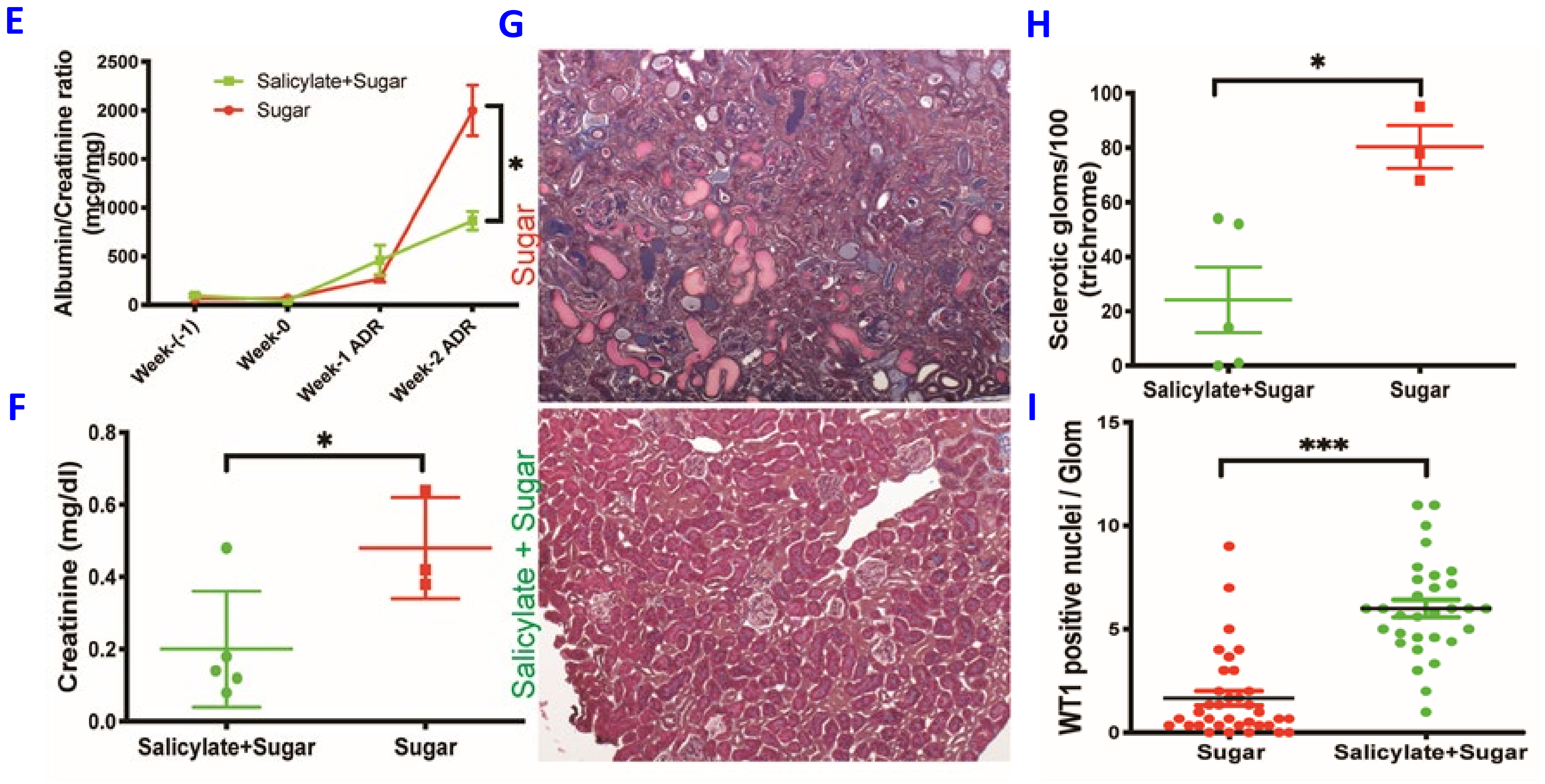


**Figure S8 (continued) E-I.** AMPK-activation confers protection from Adriamycin injury (Glomerulosclerosis): Adult BALBc mice (8-wks) were fed with 3% Sucrose solution (n=5/3) - Control Group, and with 250mg/l Sodium Salicylate in 3% Sucrose (n=5) - Experimental Group. Lines show (8E) mean Albumin:Creatinine ratio (µg/mg); dot-plots compare (8F) Creatinine (mg/dl); (8G) Representative images of MTS-stained sections shows glomerulosclerosis/collagen deposition in control group; dot-plots compare (8H) sclerosis score from Trichrome staining and (8H) mean podocyte numbers/glomerulus/animal (30 glomerular profiles/mouse) by WT1/DAPI stain; among the experimental and control group. [Line/Whiskers = Mean/SEM; unpaired t-test; *= P<0.05; MTS=Masson Trichrome stain; WT1= Wilm's Tumor-1 protein].
