## Supplemental Tables for "AMP-Kinase mediates regulation of glomerular volume and podocyte survival"

**Supplementary Tables S1-2**

**Table S1A** shows mean Vglom (μm^3^) and Vglom components (normalized to a representative control animal) by Cavalieri method in Control vs Shroom3-KD mice (>10 Glomeruli/mouse, n=8)

**Table S1B** shows mean Vglom (μm^3^) and Vglom components (normalized to a representative control animal) by Cavalieri method in Control vs Podocyte-Shroom3-KD mice (>10 Glomeruli/mouse, n=6 vs 5).

**Table S2** shows mean Vglom (μm^3^) and Vglom components (normalized to a representative control animal) by Cavalieri method in Control vs Shroom3-KD mice injected with Compound C (n=3 vs 4, respectively) used for AMPK inhibition studies.

| **Experiment Title** | **Glomerular volume**  **(Vglom x 1000μm^3^)** | | | ***Normalized volume**  **Podocyte Component** | | | **^*^Normalized volume Mesangial component** | | | **^*^Normalized volume**  **Capillary + Endothelial component** | | |
| --- | --- | --- | --- | --- | --- | --- | --- | --- | --- | --- | --- | --- |
|  | **Control**  **(Mean ±**  **SEM)** | **Shroom3-KD (Mean ±**  **SEM)** | **^**^p value** | **Control**  **(Mean ±**  **SEM)** | **Shroom3-KD (Mean ±**  **SEM)** | **^**^p value** | **Control**  **(Mean ±**  **SEM)** | **Shroom3-KD (Mean ±**  **SEM)** | **^**^p value** | **Control**  **(Mean ±**  **SEM)** | **Shroom3-KD (Mean ±**  **SEM)** | **^**^p value** |
| **TABLE S1A** | | | | | | | | | | | | |
| Shroom3-KD  (n=8 vs 8) | 167.0 ± 5.8 | 128.4 ± 4.8 | < 0.001 | 1.1 ± 0.1 | 0.9 ± 0.1 | 0.013 | 1.1 ± 0.1 | 1.0 ± 0.1 | 0.098 | 1.0 ± 0.1 | 0.9 ± 0.03 | 0.060 |
| **TABLE S1B** | | | | | | | | | | | | |
| Podocyte-Shroom3-KD  (n=6 vs 5) | 133.8 ± 6.3 | 107.8 ± 2.7 | 0.004 | 1.0 ± 0.0 | 0.8 ± 0.0 | 0.008 | 1.0 ± 0.0 | 1.0 ± 0.04 | 0.246 | 1.0 ± 0.1 | 0.8 ± 0.1 | 0.030 |
| **TABLE S2** | | | | | | | | | | | | |
| Compound C  (n=3 vs 4) | 113.7 ± 8.2 | 154.0 ± 11.3 | 0.044 | 1.0 ± 0.1 | 1.5 ± 0.0 | 0.057 | 1.0 ± 0.1 | 1.4 ± 0.3 | 0.228 | 1.0 ± 0.1 | 1.3 ± 0.1 | 0.114 |

**^*^** = Normalized to the mean of the control in each comparison

**^**^** = p values were obtained by Mann Whitney test

**Supplementary Table S3**

Table S3 showing primer pairs (with corresponding DNA sequence) used in the study.

| **S. No.** | **Primer name** | **Sequence (5' -> 3')** |
| --- | --- | --- |
| 1 | 18S rRNA-F | GGCCCTGTAATTGGAATGAGTC |
| 2 | 18S rRNA-R | CCAAGATCCAACTACGAGCTT |
| 3 | 5S rRNA-F | GGCCATACCACCCTGAACGC |
| 4 | 5S rRNA-R | CAGCACCCGGTATTCCCAGG |
| 5 | HsRPS26-F | GAAAAGAAGGAACAATGGTCGTGCC |
| 6 | HsRPS26-R | CATCGAAGACGCTCGCTTCAGAA |
| 7 | MsRPS26-F | AAGAAGAAACAACGGTCGCGC |
| 8 | MsRPS26-R | CGTCGAAGACGCTTGCTTCAGATA |
| 9 | β-actin-F | AAATCTGGCACCACACCTTC |
| 10 | β-actin-R | GGGGTGTTGAAGGTCTCAAA |
