## Supplemental Methods for "AMP-Kinase mediates regulation of glomerular volume and podocyte survival"

**Cell Culture:** Human podocyte cell line (Generous gift of Dr Moin Saleem), were expanded using RPMI-1640 (1% ITS) and DMEM (GIBCO) media, respectively. Podocytes were differentiated in Collagen coated culture plates/flasks. Protein:DNA ratio assay was performed by seeding 1000-podocytes per well in Collagen-coated 96 well plates. Briefly, cells were lysed using 1X SSC buffer, for 1 hour with intermittent shaking. DNA determination was performed as described previously (1) following addition of Hoechst 33258 (Sigma) at 1µg/ml and read at excitation wavelength 360 nm, emission wavelength 460 nm using a SpectraMax M3 microplate reader. Protein concentration was determined by Bradford protein assay (Bio Rad laboratories). Protein/DNA ratios were then calculated and shown as dot plots.

**Reverse transcription:** RNA was extracted using TRIzol and was transcribed into cDNA using a High Capacity cDNA Reverse Transcription Kit (Applied Biosystems) with starting total RNA ~ 1000 ng.

**Quantitative-PCR:** Transcript expression was assayed *in vitro/in vivo* by real-time polymerase chain reaction (qPCR) (Applied Biosystems 7500). Amplification curves were analyzed using automated 7500 software platform, via the delta-delta CT method. Human Actin was used as endogenous control. Similarly, primers were designed for human and mouse 18S RNA, 5S RNA, RPS26, Actin are listed in (Table S3).

**Western Blotting:** Cells were lysed with a buffer containing 25 mM Tris-HCl pH 7.4, 150 mM NaCl, 1 mM EDTA, 1% NP-40 and 5% glycerol, a protease inhibitor mixture and tyrosine and serine/threonine phosphorylation and phosphatase inhibitors. Lysates were subjected to immunoblot analysis using polyclonal SHROOM3 Rabbit antibody (#SAB3500818, Sigma), Fyn Rabbit polyclonal (#4023S, Cell Signaling), Actin Mouse monoclonal antibody ; (# A5441, Sigma), phospho-mTOR (Ser2448) Rabbit monoclonal antibody [EPR426(2)] (#ab109268, Abcam), mTOR (Ser2448) Rabbit polyclonal antibody (#ab2732, Abcam), phospho-AMPK (Thr172) Rabbit polyclonal antibody (#2535, Cell Signaling), AMPK Rabbit polyclonal antibody (#2532, Cell Signaling), phospho-eEF2 (Thr56) Rabbit polyclonal antibody (#2331, Cell Signaling), eEF2 Rabbit polyclonal antibody (#2332, Cell Signaling), turboGFP Mouse monoclonal antibody, clone OTI2H8 (#TA150041, CiteAb), Phospho-ULK1 (Ser555) Rabbit polyclonal Antibody (#5869, Cell Signaling), ULK1 (D8H5) Rabbit monoclonal antibody (#8054, Cell Signaling), LC3A/B Rabbit polyclonal antibody (#4108, Cell Signaling). Densitometry was performed on images of western blots using ImageJ software.

**shRNA suppression studies:** Human *SHROOM3* (2, 3) and *FYN* short hairpin clones (Dharmacon, inc, USA) were tested for optimal suppression in human podocytes. The selected GFP-tagged or (red fluorescent protein) RFP-tagged hairpins and respective scrambles were used to generate a mammalian VSV pseudotyped lentiviral expression construct. Lentiviral medium was used to infect human podocytes at 33°C. Cells were passaged in puromycin (2µg/ml)-RPMI 1640 for experiments after atleast 7-days differentiation at 37°C.

**Immunofluorescence (Podocytes):** Stably infected Shroom3-, Fyn-shRNA and Scramble-1 & 2 podocytes were plated in 12-cm wells on collagen-coated cover slips (20% rat-tail collagen, BD biosciences, San Jose CA) and allowed for differentiation for 7-10 days. Followed by formalin-fixation (4% HCHO, 0.1% TritonX100 in PBS), blocking in 3% BSA and 0.5% Triton X- 100 containing PBS. Blocked podocytes were incubated overnight with primary antibodies i.e., LC3A/B Rabbit polyclonal antibody (#4108, Cell Signaling) or LKB1 Rabbit monoclonal antibody (D60C5F10) (IHC Formulated) #13031, Cell Signaling) at a 1:100 dilution followed by 1xPBS washes and secondary antibody incubation using Alexa-Fluor 488 (Invitrogen, Life technologies, Grand Island NY) at 1:200 dilution in 3% BSA-PBS for immunofluorescence. Samples were mounted using VECTASHIELD Antifade Mounting Medium with DAPI for nuclear staining. **Immunofluorescence (Kidney Tissues):** For IF, 5-μm paraffin sections of formalin-fixed kidney-tissues were deparaffinized and processed for unmasking and antigen retrieval followed by overnight incubation with primary antibodies anti-pAMPK, anti-WT1; for fluorescence microscopy as described previously. >30 glomeruli per mouse were assayed in 40X images for pAMPK/WT-1/DAPI co-staining.

**Quantitative Image Analysis: pAMPK-Immunofluorescence:** Glomeruli were outlined using Zen pro (Zen 2.6 (blue edition)) software (40X images), and area of pAMPK-staining were measured for signal intensity and expressed as average signal intensity total area/glomerulus. All glomeruli in the biopsy tissues of the mice were included (ranging from 24-58, average of 44 glomeruli per mouse).

**Murine Shroom3 knockdown model:** Tetracycline-responsive, shRNAmir-mediated Shroom3 knockdown mouse strain based on tested shRNA guide sequences was developed with Mirimus Inc, NY. In this mouse model, Shroom3 knockdown was induced with tetracyclines by a specific shRNA hairpin in a MIR-30 backbone, selected after application of unique design algorithms(4), and followed by *in vitro* validation in murine kidney cell lines has been reported before by our group(2, 3). In the double-transgenic Chicken beta-Actin promoter (CAGS) rtTA/Nphs1-rtTA; Shroom3 RNAi mice, shRNAmir-mediated knockdown and GFP production were driven by a universal promoter and, inducible by Doxycycline feeding (DOX). Nphs1-rTTA/Shroom3 RNAi mice mice were generated for podocyte-specific shRNA expression (generous gift from Dr Miner to Dr He(5)). CAGS/Nphs1-rtTA animals were backcrossed into BALBc background. Male mice (~8-week-old) were DOX-fed for 6 weeks (600mg/g DOX chow), Non transgenic DOX-fed littermates were used as controls. Kidney tissues were collected for histology, EM, immunofluorescence (snap frozen for IF) RNA, protein. Glomeruli were extracted using DYNA-bead perfusion*.*

***For Ageing studies****,* Control and Shroom3-KD mice were aged >1 year, before DOX feeding for 6 weeks was initiated.

***AMPK inhibition studies*** were performed in control vs Shroom3-KD mice using Compound C. Adult mice (after 4 weeks of Dox feeding to induce Shroom3 knockdown) mice were injected intraperitoneally with 4 doses every 24 hours at 20mg/kg Compound C in mineral oil (vehicle) and observed till week 8 before sacrifice (15, 16). ***AMPK activation studies*** were performed in BALBc mice - FSGS model (5/6 nephrectomy) using PF-06409577. Mice were orally dosed at 50mg/kg PF-06409577 in Vehicle (0.5% methyl cellulose/0.1% Tween-80 before surgery-1 removing 2/3rd of the left kidney, followed by 100mg/kg PF-06409577 after surgery-2 and continued for 3 times a week for 6-wks (14). For histology evaluation, 85.91±26.5 glomeruli were utilized.

***Metformin protocol for aged mice:*** Metformin (dose= 300 mg/kg body weight per day(6)) in drinking water was added at week-2 of DOX feeding (after onset of proteinuria based on prior data), and continued till sacrifice.

**Uninephrectomy/5/6th Nephrectomy:** For Uninephrectomy studies, Eight-week old Doxycyline fed male control vs Shroom3-KD/ Podocyte-Shroom3-KD mice were subjected to Unilateral nephrectomy, nephrectomized kidneys (Nx-Kidney) were saved. Mice were sacrificed either 1 or 2 weeks after surgery to obtain the residual kidney (Rx-Kidney) for various morphometry, histology studies.

For 5/6 nephrectomy, 7-8-week-old mice were used. During the first surgery, 2/3rd of the left kidney was removed, by exposing it through an angular incision in the Left aspect of erector spinae muscle (left lumbar region). The upper and lower poles for 2/6 reduction of total renal mass was ligated and cut. The secured remnant 1/3rd left kidney was placed inside the muscle layer, the muscle layer and skin were then separately closed using 4-0 vicryl. After 7 days, angular Incision was made in Right aspect of erector spinae muscle (right lumbar region) exposing right kidney. 4-0 Vicryl suture was used to run a loop around the vascular pose of the kidney (hilum) (including renal artery, vein and ureter). Kidney was held with atraumtic forceps and a clean cut made lateral to the secured loop; and removed for processing and data. The secured hilum was placed inside the muscle layer. The muscle layer and skin were separately closed using 4-0 vicryl (7).

**Drug treatments:** AMPK inhibition studies were performed in control vs Shroom3-KD mice using Compound C. 7-8 week old mice (following 4 weeks of Dox feeding to induce Shroom3 knockdown) mice were injected intraperitoneally with 4 doses every 24 hours at 20mg/kg Compound C in mineral oil (vehicle) and observed till week 8 before sacrifice (8, 9).

AMPK activation studies were performed in BALB-c mice - FSGS model (5/6 nephrectomy) using PF-06409577. 7-8 week old mice were orally dosed at 50mg/kg PF-06409577 in Vehicle (0.5% methyl cellulose/0.1% Tween-80 before the first stage of surgery removing 2/3rd of the left kidney, followed by 100mg/kg PF-06409577 once one right kidney was removed and continued for 3 times a week for 42 days before sacrifice (7).

*in vitro* autophagy inhibition studies were performed in Scramble-1 and SHROOM3 shRNA podocytes using Bafilomycin A1. Bafilomycin A1 (10) was added to the culture media at Day7 of differentiation at a concentration of 100nM for 24 hours, following which the podocytes were fixed using 4% Formalin in 1xPBS and followed for immunofluorescence study (11).

**Glomerular Morphometry:**

*Tissue Processing:* Kidneys were perfused with PBS for five minutes. One-millimeter cubes were cut from the cortex and placed in glutaraldehyde. Tissue cubes were then post-fixed with 1% osmium tetroxide, dehydrated through a series of ethanol and embedded in Polybed 812 (Electron Microscopy Sciences, Hatfield, PA).

*Glomerular Volume-Weibel-Gomez Method:*  *Glomerular Volume-Weibel-Gomez Method:*  The Weibel-Gomez method was used to measure glomerular volume (Vglom) in human NS biopsies.  This method uses one PAS-stained and one Trichrome stained paraffin section from Aperio-scanned images of NS biopsies from the NEPTUNE study (12).  The areas of all complete glomerular profiles present in the section were measured by planimetry (Figure S1A).

Vglom = A^3/2^ x 1.38 µm^3^ where A is the average glomerular tuft area and 1.38 is the shape correction factor assuming glomeruli are spheres(13). The mean-Vgloms obtained from two sections within each patient were highly correlated (Fig S1B). The PAS-stained sections were used for stereological analyses.

The mean-Vgloms obtained from two sections within each patient were highly correlated (Fig S1B). The PAS-stained image data was used for clinical analyses.

*Glomerular Volume-Cavalieri Method:* Glomerular volume (Vglom) was measured using the Cavalieri Principle (14). Serial 1-µm-thick epon sections were cut using an ultramicrotome and every 10^th^ section was saved to a slide and stained with 1% toluidine blue. Using the 10x microscope objective, a map was drawn of all the glomerular profiles present in the first section. Using the map and the subsequent sections, newly appearing glomeruli were mapped and sequentially numbered. Only complete glomeruli defined as having a section before its appearance on a section and after its disappearance were used for measurements (Figure S1A). For each complete glomerulus, the 100x objective was used to image each profile from the individual glomeruli. An average of 6.9 profiles was imaged per glomerulus. Using Adobe Photoshop’s layers function a grid of points (100mm apart) was superimposed over each profile and the number of grid points “falling” over a profile was counted. V_glom_ = 10 x (d/mag)^2^ x ∑Points µm^3^ where 10 is the distance between profiles in µm, d is the distance between grid points in µm, mag is the magnification and ∑Points is the sum of grid points “falling” on the profiles from a glomerulus (Figure S1A). An average of 11.0 glomeruli per animal was measured and an average of 1509 grid points were counted over all the glomeruli per animal.

*Glomerular Component Volume:* One-half of the images obtained for Cavalieri were used for estimation of the volumes of glomerular components (Fig S1B-D). For odd numbered glomeruli the odd numbered images were used and for even numbered glomeruli the even numbered images were used. A representative image used for glomerular component volume analyses is shown in Fig S1B. Glomerular profile was first defined by a minimal polygon around the glomerular profile in image. Podocyte substructure such as pseudocysts are visible on light microscopy using this technique as published earlier(15). As described previously(2, 16), we defined four glomerular components: podocyte, capillary lumen + endothelial cell, mesangium, and “other” (pseudo-color in Fig S1C). The “other” component was defined as Bowman’s space, glomerular basement membrane and non-resolvable areas within the glomerular profile. The “other” data was not analyzed in this study. Using Photoshop’s layers function a grid of points was superimposed over the images. The number of points falling on each component was counted (Fig S1D). The volume fraction of component X per glomerulus [V_v_(Comp X/glom)] = ∑Points_Comp X_ /∑Points_Total_ µm^3^/µm^3^ where ∑Points_Comp X_ is the number of points “falling” on component X, ∑Points_Total_ is the number of points “falling” on all four components of the glomerulus(13). An average of 169.2, 152.4, 255.6, and 78.6 points fell on podocyte, mesangium, capillary lumen + endothelial cell and “other” respectively. The volume of a Component X (CompxVglom) is calculated by multiplying the component volume fraction by the glomerular volume: CompxVglom = V_v_(Comp X/glom) x Vglom	µm^3^.

*Podocyte Number:* The fractionator/disector method was used to count the number of podocytes per glomerulus (N_podo_) (17, 18). Podocyte nuclei were surrogates for podocytes assuming only one nucleus per podocyte. For this method pairs of sections are needed. We used the 1-µm epon sections available for the Cavalieri determination of Vglom. In addition, at the time of saving sections for the Cavalieri method the section adjacent to the Cavalieri section was saved to make disector pairs. At the time of imaging for the Cavalieri method the adjacent section was also imaged. Using Photoshop’s pencil tool all the podocyte nuclei profiles were mark on all the sections. Then looking at the pairs of images next to each other the number of podocyte nuclei profiles from nuclei present in one section but not in the other were counted. This was repeated for each disector pair from a glomerulus. An average of 6.9 disector pairs per glomerulus was available. N_podo_ = ∑Q^-^ x 10, where ∑Q^-^ is the number of nuclei profiles from podocytes present in one section but not present in the other section of a disector pair and 10 is the reciprocal of the fraction of the glomerulus sampled. An average of 128.8 Q^-^ were counted per kidney.

**Statistical analysis:** De-identified clinical and demographic information was obtained for the NS morphometry cohort and linked to morphometry measurements using unique-IDs. ***For human data,*** univariate comparisons of clinical factors and demographics between NS categories were done using ANOVA (Kruskal Wallis for corresponding nonparametric analysis with post-test Dunn’s test) for continuous variables, and Chi-Square for proportions. Spearman correlation coefficient was used to compare Vgloms in two random sections within the same patient. Cox proportional hazard models were used for multivariable survival associations, including clinic-demographics identified as significantly different in uni-variable analyses. NEPTUNE determined outcomes of End stage renal failure, eGFR decline ≥ 40% from baseline, or a composite of these events were evaluated as outcomes. Time from biopsy to event was utilized. ***For in vitro and in vivo experiments,*** unpaired t test was used to analyze data between two groups. *in vitro* experiments were repeated multiple times to obtain standard deviations, and representative experiments are shown. Univariate comparisons of continuous variables were done using unpaired t-test (Mann-Whitney test for corresponding non-parametric analysis). When >2 groups were compared, ANOVA or Kruskal Wallis (for non-parametric analyses with post-test Dunn’s test) was used. Statistical significance was considered with two-tailed P<0.05. ***Software:*** Graphpad Prism Version 9 (Graphpad, LaJolla, CA) and SPSS version 24 (IBM, NY) were used for analyses.

**Study approval:** Institutional IACUC approved protocol was available for all mouse experiments performed according to humane endpoints. The human data in this work was obtained via a NEPTUNE anciliary study to examine morphometric, genomic and signaling changes in MCD, FSGS and all NS. The anciliary study was approved in 2018 and renewed in 2020 to permit analyses. Institutional IRB approval (Exemption 4) was obtained to examine de-identified NEPTUNE data.

**Supplemental References:**

1.	Rao J, and Otto WR. Fluorimetric DNA assay for cell growth estimation. *Anal Biochem.* 1992;207(1):186-92.

2.	Wei C, Banu K, Garzon F, Basgen JM, Philippe N, Yi Z, et al. SHROOM3-FYN Interaction Regulates Nephrin Phosphorylation and Affects Albuminuria in Allografts. *J Am Soc Nephrol.* 2018;29(11):2641-57.

3.	Menon MC, Chuang PY, Li Z, Wei C, Zhang W, Luan Y, et al. Intronic locus determines SHROOM3 expression and potentiates renal allograft fibrosis. *J Clin Invest.* 2015;125(1):208-21.

4.	Premsrirut PK, Dow LE, Kim SY, Camiolo M, Malone CD, Miething C, et al. A rapid and scalable system for studying gene function in mice using conditional RNA interference. *Cell.* 2011;145(1):145-58.

5.	Lin X, Suh JH, Go G, and Miner JH. Feasibility of repairing glomerular basement membrane defects in Alport syndrome. *J Am Soc Nephrol.* 2014;25(4):687-92.

6.	Bachmanov AA, Reed DR, Beauchamp GK, and Tordoff MG. Food intake, water intake, and drinking spout side preference of 28 mouse strains. *Behav Genet.* 2002;32(6):435-43.

7.	Kir S, Komaba H, Garcia AP, Economopoulos KP, Liu W, Lanske B, et al. PTH/PTHrP Receptor Mediates Cachexia in Models of Kidney Failure and Cancer. *Cell Metab.* 2016;23(2):315-23.

8.	McCullough LD, Zeng Z, Li H, Landree LE, McFadden J, and Ronnett GV. Pharmacological inhibition of AMP-activated protein kinase provides neuroprotection in stroke. *J Biol Chem.* 2005;280(21):20493-502.

9.	Abdulrahman RM, Boon MR, Sips HC, Guigas B, Rensen PC, Smit JW, et al. Impact of Metformin and compound C on NIS expression and iodine uptake in vitro and in vivo: a role for CRE in AMPK modulation of thyroid function. *Thyroid.* 2014;24(1):78-87.

10.	Riediger F, Quack I, Qadri F, Hartleben B, Park JK, Potthoff SA, et al. Prorenin receptor is essential for podocyte autophagy and survival. *J Am Soc Nephrol.* 2011;22(12):2193-202.

11.	Bhaskar Das DPW. LKB1-AMPK ACTIVATORS FOR THERAPEUTIC USE IN POLYCYSTIC KIDNEY DISEASE. <https://pubchem.ncbi.nlm.nih.gov/patent/US2017334892#section=Patent-Submission-Date>.

12.	Lemley KV, Bagnasco SM, Nast CC, Barisoni L, Conway CM, Hewitt SM, et al. Morphometry Predicts Early GFR Change in Primary Proteinuric Glomerulopathies: A Longitudinal Cohort Study Using Generalized Estimating Equations. *PLoS One.* 2016;11(6):e0157148.

13.	ER W. *Stereological Methods*. London: Academic Press; 1979:44-5.

14.	Gundersen HJ, and Jensen EB. The efficiency of systematic sampling in stereology and its prediction. *J Microsc.* 1987;147(Pt 3):229-63.

15.	Zhou Y, Castonguay P, Sidhom EH, Clark AR, Dvela-Levitt M, Kim S, et al. A small-molecule inhibitor of TRPC5 ion channels suppresses progressive kidney disease in animal models. *Science.* 2017;358(6368):1332-6.

16.	Basgen JM, and Sobin C. Early chronic low-level lead exposure produces glomerular hypertrophy in young C57BL/6J mice. *Toxicol Lett.* 2014;225(1):48-56.

17.	Sterio DC. The unbiased estimation of number and sizes of arbitrary particles using the disector. *J Microsc.* 1984;134(Pt 2):127-36.

18.	Bai XY, and Basgen JM. Podocyte number in the maturing rat kidney. *Am J Nephrol.* 2011;33(1):91-6.
